# 15 years of introgression studies: quantifying gene flow across Eukaryotes

**DOI:** 10.1101/2021.06.15.448399

**Authors:** Andrius J. Dagilis, David Peede, Jenn M. Coughlan, Gaston I. Jofre, Emmanuel R. R. D’Agostino, Heidi Mavengere, Alexander D. Tate, Daniel R. Matute

## Abstract

With the rise of affordable next generation sequencing technology, introgression — or the exchange of genetic materials between taxa — is widely perceived to be a ubiquitous phenomenon in nature. Although this claim is supported by several keystone studies, no thorough assessment on the frequency of introgression in nature has been performed to date. In this manuscript, we aim to address this knowledge gap by providing a meta-analysis of the most comprehensive survey of introgression studies in Eukaryotes to date (724 papers with claims of introgression). We first examined the evidence given to support introgression, and if/how the lines of evidence have changed across time. We then collated a single statistic, Patterson’s *D*, that quantifies the strength of introgression across 123 studies to further assess how taxonomic group, divergence time, and aspects of life history influence introgression. We find three main results. Studies on introgression are much more frequent in plants and mammals than any other taxonomic group. The study of introgression has shifted from a largely qualitative assessment of whether introgression happens, to a focus on when and how much introgression has occurred across taxa. The most often used introgression statistic, Patterson’s *D*, shows several intriguing patterns suggesting introgression reports may be biased by both differences in reporting criteria and sequencing technology, but may also differ across taxonomic systems and throughout the process of speciation. Together, these results suggest the need for a unified approach to quantifying introgression in natural communities, and highlight important areas of future research that can be better assessed once this unified approach is met.

## INTRODUCTION

Genome sequencing has revealed that instances of hybridization and introgression — transfer of alleles from one species into a different one — are not rare in nature. Introgression can have myriad effects, and though it is most commonly thought to be deleterious, introgression may also provide the raw genetic materials for adaptation and speciation (Heiser 1973; Rieseberg and Wendel 1993; Dowling and Secor 1997; Arnold and Martin 2009; Suarez-Gonzalez, et al. 2018). Examples ranging from disease vectors (Lee, et al. 2013; Fontaine, et al. 2015; Norris, et al. 2015) to humans have revealed that allele transfer can be instrumental for range expansion, adaptation and even speciation. For example, the *EPAS*1 allele responsible for Tibetan high altitude adaptation most likely introgressed from Denisovan populations (Huerta-Sánchez, et al. 2014; Racimo, et al. 2015). On the other hand, introgressed genes may bear certain costs (Harris and Nielsen 2016) - *Neanderthal* variants in human populations have been associated with high health risk for SARS-CoV-2 infections (Zeberg and Paabo 2020, 2021). However, the relative importance of introgression for adaptation remains largely unknown, mainly because the commonality of introgression across species also remains unknown.

The susceptibility of genomes to introgression has historically been a subject of lively debate among evolutionary biologists (Barton 2001; Mallet 2005; Schwenk, et al. 2008; Payseur and Rieseberg 2016). While classically controversial, there is now general consensus among evolutionary biologists that introgression can occur between species; however, the frequency with which introgression occurs and the genomic and environmental conditions that facilitate or preclude gene exchange between species are relatively unresolved. Nonetheless, there are good reasons to believe introgression may vary in frequency across the tree of life as well as over the course of speciation, as illustrated by examining the conditions that must be met for introgression to occur. We discuss each of these in turn.

For introgression to take place, hybrids must first form, and then be able to serve as a bridge for genetic material to cross species boundaries. Thus, introgression requires at least a degree of sympatry and incomplete prezygotic isolation. As a result, taxa with larger ranges or weaker mate choice are expected to show higher rates of introgression. Furthermore, the hybrids must be viable at least to the age of reproduction and be partially fertile to produce advanced backcrosses. While hybrid fitness is expected to decrease as species continue to diverge (Prager and Wilson 1975; Coyne and Orr 1989; Coughlan and Matute 2020; Satokangas, et al. 2020), it is possible that introgression occurs rather freely until a critical threshold of low fitness in hybrids is developed (Barton 2001; Roux, et al. 2016). Since the rate at which reproductive isolation evolves differs widely by taxa (Coughlan and Matute 2020), there may likewise be differences in the degree of introgression between species. Several historic reviews have examined the frequency of hybridization in general (Knobloch 1972; Dowling and Secor 1997; Payseur and Rieseberg 2016), but none to our knowledge have examined introgression specifically.

Nonetheless, the production of advanced intercrosses is not a guarantee that the introgressed material will remain in the recipient species. While it is true that selection might increase the frequency of alleles important for adaptation, selection might also reduce the frequency of alleles that diminish the hybrid fitness (i.e., alleles involved in hybrid incompatibilities or deleterious alleles that contribute to hybridization load) (Harris and Nielsen 2016; Martin and Jiggins 2017). Theoretical and simulation work suggests that the linkage between positively and negatively selected alleles is crucial to determine the fate of an introgressed allele (Barton and Hewitt 1989; Liang and Nielsen 2014; Shchur, et al. 2020). Three factors play an important role. The timing of introgression determines the likelihood of encountering an introgressed allele. Alleles that are selected against (and to a lesser extent neutral), will only persist in a population if admixture is recent (Racimo, et al. 2015; Harris and Nielsen 2016). Similarly, one will observe more negatively selected alleles in instances in which admixture is continuous than in incidences with a single pulse of admixture because the input of introgressed materials (including potentially deleterious alleles) will be continuous. Here, again, taxa with higher degrees of sympatry are expected to experience increased rates of introgression. On the other hand, reproductive isolation accumulates faster in sympatric species pairs (Coyne and Orr 1989; Matute and Cooper 2021), and introgression may be limited solely to species in secondary contact after a period in allopatry. In cases in which multiple alleles affect fitness, the genetic distance between two alleles, which ultimately depends on the recombination landscape across the genome, will also define the introgression of both positive and negative variants. These factors can therefore create differences in introgression not only between taxa but also systematically within different parts of the genome. It has been observed, for example, that signals of introgression are stronger in regions of higher recombination (Brandvain, et al. 2014; Sankararaman, et al. 2014; Harris and Nielsen 2016; Juric, et al. 2016; Muirhead and Presgraves 2016; Edelman, et al. 2019; Petr, et al. 2019). The evolutionary determinants of the fate of introgression are an active area of research.

While individual studies across focal taxa have been instrumental in revealing specific instances of introgression, the relative occurrence of introgression across taxa remains unknown. To address the differences in introgression across taxa, a comparative approach that consolidates measurements of introgression is required. The probability of ongoing migration has been elegantly analyzed for some taxa by Roux, et al. (2016), but aside from a few general reviews pointing to the increasing evidence for introgression (Payseur and Rieseberg 2016), no systematic analysis of introgression has been performed across multiple kingdoms of eukaryotes. The difficulty, in part, has been in quantifying introgression - while shared haplotypes or reduced divergence within a particular region are evidence for potential introgression between two species, they are difficult to compare between species. As researchers moved from sequencing individual genes to entire genomes, novel methods to quantify the degree of introgression have been developed. One of the earliest and most successful is Patterson’s *D* (Green, et al. 2010; Durand, et al. 2011), which has spawned a series of other so-called *f-*statistics (see Table 1). These statistics evaluate the degree to which gene frequencies or tree topology patterns support introgression versus incomplete lineage sorting (Supplementary Figure 1). While care must be applied when evaluating any of the *f-*statistics, they represent an opportunity to compare the frequency and strength of evidence for introgression across different taxa. Ideally, *f-*statistics would be computed for a variety of taxa using a single set of approaches, as has been done by Hamlin, et al. (2020), but it is difficult to scale this approach using comparable data across eukaryotic life. Alternatively, published data can be used to investigate differences in introgression across taxa. In this manuscript, we undertake the latter approach.

**Table 1.**
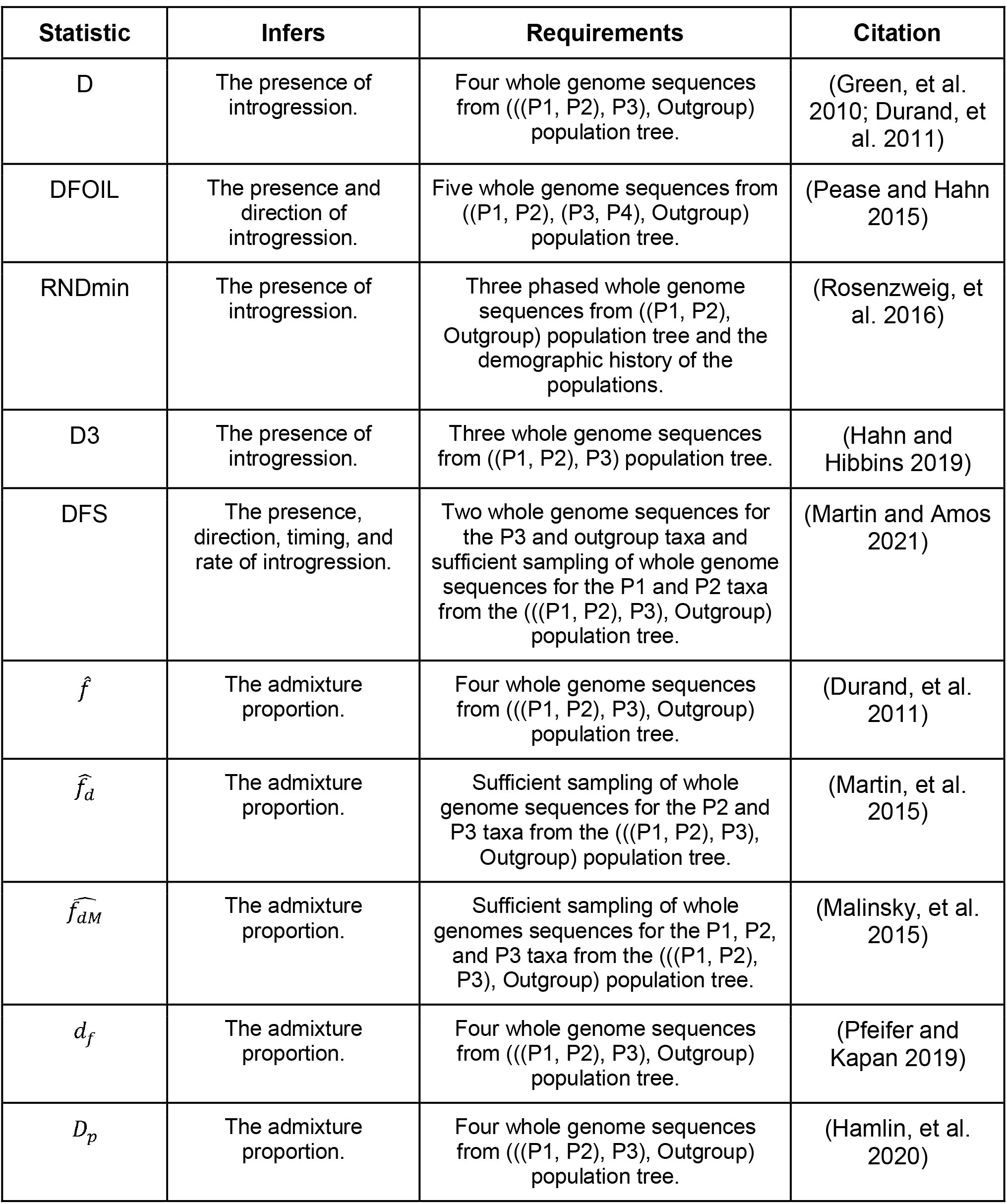
List of introgression summary statistics that were binned into f-statistics.

By searching through 724 studies published since 2005 with claims of introgression, we were able to evaluate the evidence for introgression across time and diverse eukaryotic taxa. When *f*-statistics were available, they were extracted, resulting in a dataset of nearly 20,000 Patterson’s *D* values between more than 1,000 species pairs. The resulting dataset was used to ask whether there were differences in introgression between taxa, and how evidence for introgression is impacted by sequencing technology, genetic divergence and several life-history traits. While we identify several intriguing patterns, our meta-analysis exposes the need for clearer reporting criteria for introgression studies, as well as further efforts at comparative work in introgression.

## RESULTS

We first identified 1,889 papers that either fell into our Web of Science query (“Introgression (AND) genome”) or cited one of several methods papers for *f-*statistics (Green, et al. 2010; Martin, et al. 2015). Papers were then manually evaluated for claims of introgression, resulting in 724 papers with claims of introgression between 2005 and February 2021. Papers were annotated for the biological system, the evidence presented, and for the data types used (Supplementary File 1). This survey revealed changes in three different aspects regarding the methodology of how introgression has been historically studied: (1) the systems used, (2) the genomic data used, and (3) the methods used to first identify, and then quantify introgression. We discuss these major trends in studies of introgression in the following sections.

### Introgression has been mostly studied in plants and mammals

Our data compilation revealed that introgression has historically been studied primarily in two classes — flowering plants and mammals (Figure 1A). Historically, hybridization and introgression have been seen as stronger drivers of evolution in plant biology than other fields (Anderson and Stebbins Jr 1954; Knobloch 1972; Heiser 1973; Rieseberg and Wendel 1993; Rieseberg 1997). Another potential reason for the focus on plants and mammals might be the interest in the role of introgression in domesticated taxa – some of the earliest well studied cases of introgression come from studies of domesticated taxa (Ellstrand, et al. 1999; Grabenstein and Taylor 2018; Ottenburghs 2021). While the relative frequency of plant and mammal studies has decreased over time, this is not caused by a decrease in the interest in these groups. The absolute frequency of studies in introgression in the group (i.e., number of total studies per year) has actually increased over time (Figure 1A) indicating that the decreased relative frequency is caused by an increase in the number of introgression studies in other taxa.

**Figure 1:**
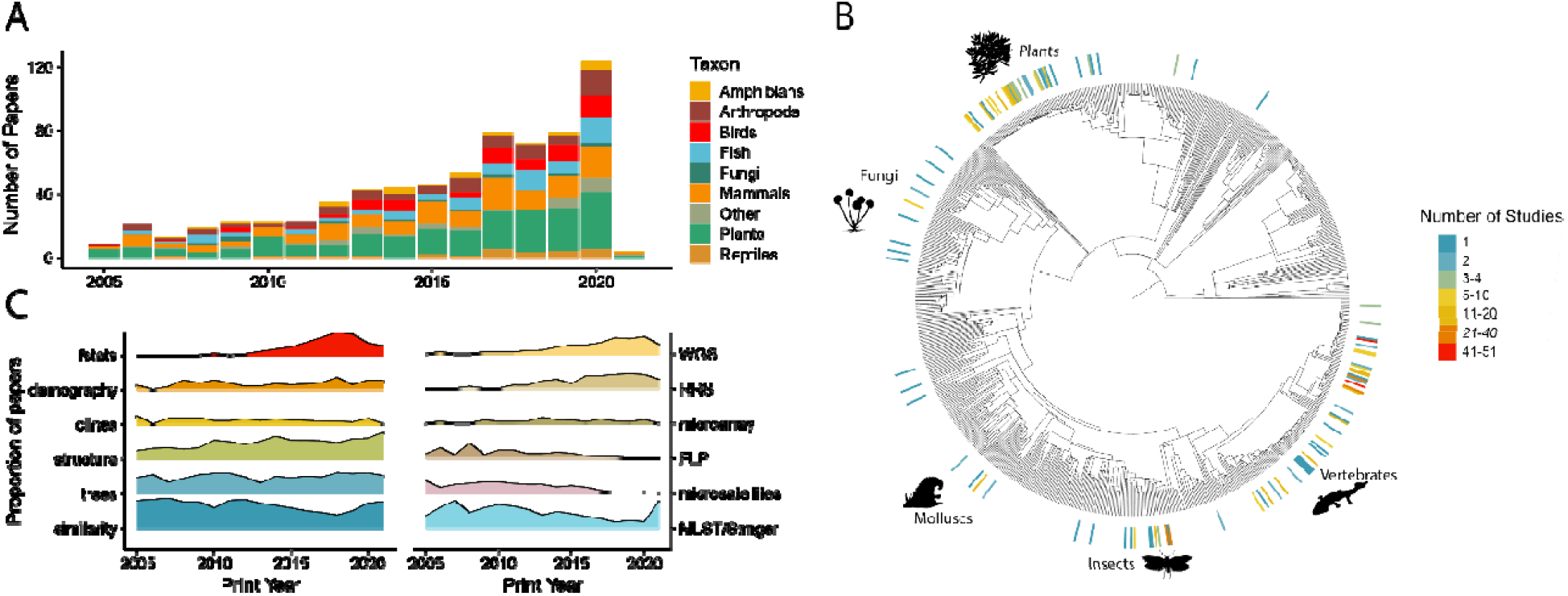
Frequency of introgression studies since 2005 across fields (A), with the number of papers per eukaryotic order shown in (B). Changes in the statistics used to identify introgression (C, left panel) have accompanied changes in sequencing technology (C, right panel).

The slow increase in new systems with evidence for introgression has resulted in at least a few case studies across most major lineages of Eukaryotes (Figure 1B). While we only identified a few studies each in mollusks, flatworms, and fungi other than *Saccharomyces*, many of these studies occurred in the last 5 years (Supplementary Figure 2). Taxonomic orders with the largest number of studies also represent a fairly broad sampling of eukaryotes, with representatives of mammals, birds, plants, insects and amphibians appearing in the top 10 orders by numbers of studies (Supplementary Table 1).

In spite of these expanded taxonomic interests, large sections of the tree of life remain largely unexplored for the occurrence of introgression. Our survey failed to identify a single study in protists, and found only a single study in algae, for example. Non-insect invertebrates and fungi, in particular, have seen few studies examining introgression given they represent a vast amount of biological and phylogenetic diversity. Given the extensive genome resources available for fungi (Stajich, et al. 2012; Matute and Sepúlveda 2019; James, et al. 2020; Li, et al. 2021) and the abundance of verbal arguments that suggest that hybridization is one of the leading forces in fungal evolution (Nelson 1963; Steensels, et al. 2021), the time is ripe to determine whether gene exchange is more prevalent in fungi than in other clades. Even within insects, most studies have occurred either in Dipterans or Lepidopterans, with few studies in other insect systems. In Coleopterans, for example, thought to be among the most speciose groups, only 3 studies of introgression filled our criteria. While it is clear that introgression seems to be common, it is too early to assess whether it is ubiquitous across eukaryotic species. Studies in non-model systems are therefore needed to fully quantify the prevalence of introgression across Eukaryotes.

### Larger numbers of loci are used to identify introgression

The study of introgression has experienced a transformation over time. Historically, introgression was inferred from morphological data (Anderson and Hubricht 1938). This approach would leverage the observation that hybrids would sometimes show intermediate trait values and mismatch of diagnostic traits but would necessarily conflate phenotypic variability, cryptic species boundaries, and true hybridization (Heiser 1973). Yet, this approach led to the assertions that hybridization was an important driver of evolution across multiple taxa (Anderson and Stebbins Jr 1954; Lewontin and Birch 1966).

The advent of DNA typing and sequencing were then incorporated to study the magnitude of gene exchange between species, the focus of this piece. Molecular tools were readily incorporated into the detection of hybrids and introgression historically (see Avise (2004), Rieseberg and Wendel (1993) for reviews). Across our dataset, we find six types of DNA data that have been used to infer gene exchange: Figure 1C shows how different data collection approaches have changed to incorporate information from multiple papers into synthetic pieces. Multi-locus sequencing typing (MLST)/Sanger based methods, in which a small region of DNA is directly sequenced, dominated the early 2000s. While this approach allowed for direct comparison as homology was almost always clear (with the exception of gene duplications), they were often cumbersome and slow as they required amplifying, cloning, and sequencing individual loci for each sample. For that reason, the number of loci was usually restricted to a handful (fungal studies typically used 1-15 loci to define species, for example (Matute and Sepúlveda 2019)).

Methods that captured larger numbers of loci, such as restriction/amplified fragment length polymorphism sequencing (FLP category in Figure 1) as well as microsatellite-based approaches complemented MLST data, but never fully replaced it. These methods increased in popularity because they were cheap, easy to implement, and were likely to find polymorphic sites. Unlike MLST methods, determining homology was a challenge in FLP and microsatellites studies, a fact which would prevent incorporating information from multiple papers into synthetic pieces. Additionally, at least for microsatellites, the high levels of homoplasy in the form of retromutation would create difficulties in determining when a potential case of hybridization was instead caused by back mutation (Putman and Carbone 2014).

While these approaches to sample the genome suggested that hybridization could indeed occur in nature, they were limited in their scope as they were more likely to reveal recent instances more than multigenerational admixture events. Since they sample such a small portion of the genome they could not reveal the broad patterns of diversity that interspecific crosses would leave in the genome. Microarrays (SnpChip, GoldenGate arrays, Beadchip, etc.) allow for larger sampling across the genome, but are only practical in taxa in which the cost of developing a micro-array could be offset by long term usage (primarily model systems). Next generation genome sequencing allowed even non-model taxa to be studied in greater detail. These methods of either whole genome sequencing (WGS) or reduced representation sequencing (RRS — including restriction-site associated DNA (RAD), genotype-by-sequencing (GBS), transcriptome and exome methods) rapidly replaced all other categories, and coincided with growing numbers of studies of introgression outside plants and mammals. The rise of whole genome sequencing also accompanied a shift in how introgression was identified. The ability to sample the entire genome (or at least a sizable fraction of the genome) gave researchers the newfound opportunity to describe and quantify global measures of introgression as well as identify putatively introgressed regions. This “genomic revolution” also ushered in many new methods that take advantage of having genetic information for more than a handful of loci, and allowed for a paradigm-shift in the field’s perspective of introgression from “Does introgression occur?” to “How often and where does introgression occur?”.

### Modern methods allow not just identification, but quantification, of introgression

The approaches used to study introgression have evolved alongside the expanding sequencing methods available to researchers (Figure 1C). While a cottage industry of developing new methods to detect introgression has arisen in the last decade (Wangkumhang and Hellenthal 2018; Hibbins and Hahn 2021), we found in practice only a subset of methods were used. To describe the usage of each of the methods to infer introgression, we binned the type of evidence to detect introgression in six different categories (Figure 1C). The first one, “Sequence similarity”, included any evidence based on direct sequence comparisons, including relative and absolute metrics of differentiation (*F*_ST_, *d*_xy_), and haplotype sharing. The benefits and weaknesses of these approaches to detect introgression have been discussed elsewhere (Smith and Kronforst 2013; Martin, et al. 2015; Hibbins and Hahn 2021). Due to their simplicity, these methods are a frequent feature of introgression studies, even if they do not provide definitive evidence of introgression. A second family of analyses uses clinal changes along space or along the genome (“cline’’ group in Figure 1C). These methods examine how ancestry at a single locus changes along either geographic distance or a set of populations with varying levels of admixture. Loci whose clines deviate significantly from the genome average are potentially adaptively introgressing (Barton and Gale 1993; Fitzpatrick 2013; Jofre and Rosenthal 2021). Clinal approaches require sampling in many populations, and so have historically been rarer than other approaches (Figure 1C). Third, tests can leverage tree topology; these analyses can use information from just a pair of genes or more modern methods like TreeMix (Pickrell and Pritchard 2012) or QuIBL (Edelman, et al. 2019) (“tree” group in Figure 1C). Inferred gene tree topology as well as branch lengths can be used to infer both the presence and sometimes timing of introgression, making phylogenomic methods both powerful and popular (Hibbins and Hahn 2021). While the way trees are used to detect introgression has changed with the addition of novel methods, they have remained popular in studies of introgression throughout the period covered herein. Fourth, clustering analyses such as STRUCTURE (Pritchard, et al. 2000) use allele frequency differences to arrange samples into the most likely clustering; these methods have been expanded to infer the precise contribution of the clusters into admixed genomes (e.g., fastSTRUCTURE (Raj, et al. 2014) or ADMIXTURE (Alexander, et al. 2009; Alexander and Lange 2011)). Clustering based methods are intuitive, do not require vast amounts of data, and so stayed popular throughout our study period. The fifth major group represents evidence through demographic model fitting. By fitting explicit demographic models with migration, researchers can identify the timing and/or the frequency of introgression events. As these methods work best with larger amounts of data, are computationally complex and require a baseline knowledge of the evolutionary history of the group, they have remained relatively unpopular in the studies in our data-set throughout the years (Figure 1C). Finally, Patterson’s *D* and all similar statistics were binned into the *f*-statistic category. These statistics compare the likelihood of incomplete lineage sorting and introgression to determine the strength of evidence for introgression (Supplementary Figure 1). Comparing the prevalence of the six groups, methodology to detect introgression has remained relatively constant, except for the addition of *f*-statistics in the 2010s.

Before the ubiquity of next generation sequencing approaches, studies were limited by the amount of genomic data they had available and thus the most frequent evidence fell into either sequence similarity or tree-based approaches. As larger numbers of individuals began to be sequenced, clustering based methods rapidly increased in popularity, becoming present in over half the papers identified in this study from the mid 2010s. While these methods may sometimes be used to infer admixture proportions, they are more often used simply to test for the presence of introgression. Since the publishing of Green, et al. (2010), *f*-statistics, which can be used to estimate admixture proportion, have exploded in popularity and are now found in more than half of any publication with evidence for introgression (Figure 1C, red area). These methods are intractable without sampling large numbers of loci, and indeed we find a significant correlation between whole genome sequencing and the use of *f*-statistics as well as demographic model fitting (Supplementary Figure 2). While we did not parse out how granular each study’s conclusions were, modern approaches like 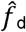 allow researchers to not only ask how much of the genome shows evidence for introgression, but also which particular parts of the genome show this signal (Martin, et al. 2015; Martin and Jiggins 2017; Hibbins and Hahn 2021). The field has therefore experienced a shift from simply finding evidence for introgression, to quantifying it, and will likely shift to examining both the timing and exact genes crossing species boundaries in the coming years.

While evidence for introgression has increased across eukaryotes, it is still unclear whether it occurs more frequently in some taxa than others. The variety of methodologies to identify introgression presents some barriers for comparative analyses — it is difficult to quantify the strength of evidence for introgression between two STRUCTURE studies, for example, and practically impossible to compare introgression between a study using tree-based methods versus demographic modeling. Since *f*-statistics provide a unified metric to measure the magnitude of introgression between species, we focused on this metric for the rest of the analyses. We were able to extract *f*-statistics from a total of 208 papers (Supplementary File 2). For each paper, we extracted the identity of the focal introgressing species, the *f*-statistic, and its significance. Many of the recently developed *f-*statistics are potentially more robust to demographic noise and use of wrong tree topologies than the original Patterson’s *D* (Hibbins and Hahn 2021). In practice, Patterson’s *D* values represented the vast majority of the data with over 30,000 Patterson’s *D* values from 123 studies. The next most frequent statistic, *f*_4_, was only available for 5 studies, although it is worth noting that Patterson’s *D* represents a specific configuration of the *f*_4_ statistic. 419 values of the _D_ statistic across 14 papers were also obtained, again representing a small number of taxa. As a result, we focused primarily on results from Patterson’s *D* statistic, as they presented the most potential comparisons between taxa.

Since Patterson’s *D* provides a common framework to determine the relative frequency of introgressed alleles in a genome, we used this metric to test three different hypotheses. First, we assessed whether introgression varies across different taxonomic groups. Second, we determined whether there was a decrease in the evidence for introgression as the divergence age of the parental species increased. Finally, we study whether human association affects introgression across our entire dataset, and whether life history traits affect the likelihood of introgression in plants. We describe the results for these tests in the paragraphs that follow.

### Introgression may be stronger in plants than animals or fungi

One of the oldest debates in speciation genetics is whether plants and animals differ in their propensity to produce hybrids (Dowling and Secor 1997; Chen, et al. 2018). This argument can be extended to a more inclusive taxonomic base: does the amount of introgression differ across taxonomic groups? To address this question, we fit a mixed model to determine whether different taxonomic groups showed differences in the amount of introgression as detected by Patterson’s *D*. To control for differences in study approaches (in terms of sequencing technology, differences in data filtering, and which values of Patterson’s *D* were reported) we used two different sets of random effects. In the most conservative of our models, both study and introgressing species pair were considered random effects. While pair identity did not explain a large amount of variation (3.4-28% of variance explained, Supplementary Table 1), reference consistently explained a large amount of variance (4-59%, Supplementary Table 1). However, since each study will only represent species within the same class, and often the same order and genus, this approach may assign variation due to biological differences to random study effects. Thus, our second approach includes species pair, genomic data type and reporting criteria (only significant values, all possible values, or a focal subset of values) as random effects. Both of these effects explained enough variation to be retained in our mixed model (Supplementary Table 1). We focused on the kingdom, phylum and class categories for fixed effects on reported Patterson’s *D*. In total we fit 30 linear models to study the differences in introgression between taxa. Table 2 and Supplementary Tables 2-9 show the results for these linear models, with models using different subsets of data separated into groups. Across a variety of models, with varying filtering for significance and quality, models which include effects of phylogenetic kingdom and class were preferred to those that did not. While the improvement in AIC for these models was modest, we performed a post-hoc ANOVA to test for significant factors and used Least-Squares Means to obtain pairwise differences in marginal means between different taxa. On the kingdom level, significant differences are only found between animals and plants (Supplementary Tables 4-7). Class level differences are driven by Polypodiopsida (ferns) and Pinopsida (conifers) — two plant classes with Patterson’s *D* values from only a single study each (Supplementary Tables 4-7). In less conservative versions of the models, differences between Actinopteri (ray-finned fishes minus bichirs) and other classes are also sometimes significant (Supplementary Tables 5,7). To further minimize the effect of individual studies, we then excluded classes with fewer than two studies from the analysis, finding significant differences between Fungi and Metazoa at the kingdom level, and Actinotperi and 5 other classes, as well as Mammalia and Sordariomycetes (Supplementary Table S8).

**Table 2:**
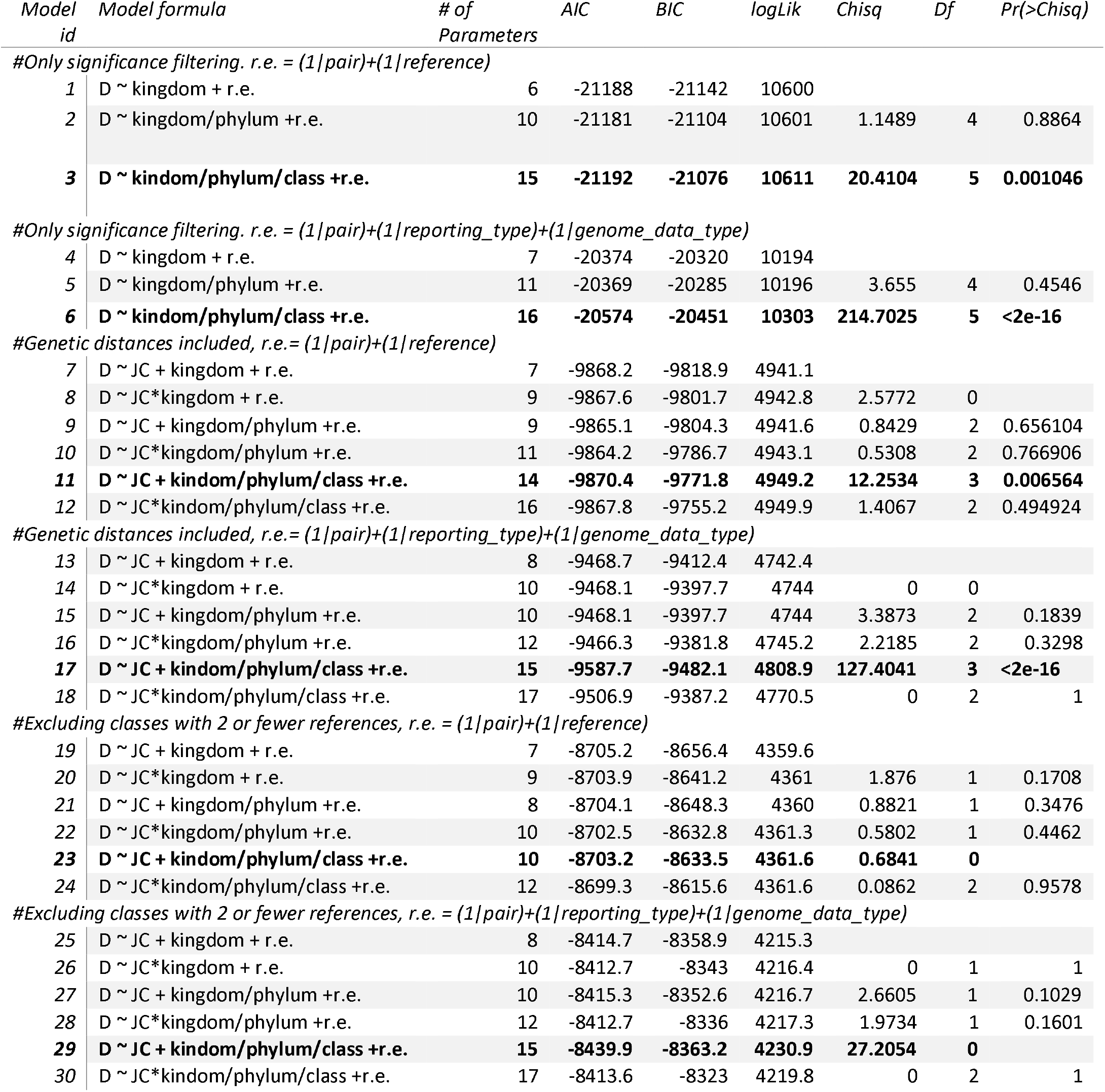
Mixed model selection for various data subsets. Best models for each category were selected by minimizing AIC.

### Evidence for introgression is weaker between more diverged species

One of the expectations of hybridization is that as divergence increases between the parental species, the number of incompatibilities increases at a fast pace (Satokangas, et al. 2020). Since the amount of residual introgression after hybridization has been hypothesized to be affected by the density of hybrid incompatibilities (Veller, et al. 2019), then hybridization between more divergent species should lead to lower signals of admixture (Wiens, et al. 2006; Hamlin, et al. 2020). We next annotated our dataset with genetic distances between pairs of species showing evidence for introgression. For each introgressing species pair, we searched for any sequences in the NCBI nucleotide database (2018) and obtained the Jukes-Cantor distance (Jukes and Cantor 1969) between reciprocal best BLAST hits. In this fashion we annotated 9,804 values of Patterson’s *D* with genetic distance. We then fit mixed models including genetic distance as a predictor of Patterson’s *D*. First, we tested whether there was an effect of genetic distances across the whole dataset. We found that genetic distance does have a significant relationship with Patterson’s *D* (slope = -0.073, SE = 0.0335, p = 0.029) when all the data are considered together (Supplementary Table 3). This relationship varies slightly among our best fit models, ranging between a slope of -0.15 to -0.06; all but one model showed that Patterson’s *D* was negatively associated with genetic distance. Including taxon specific slopes at the phylum or class level did not improve the models (as determined by AIC comparisons, Table 2). The kingdom and class effects were similar in their effect size to those found in the complete dataset without genetic distance.

Next, we explored whether there was heterogeneity in the relationship between the amount of introgression and the genetic distance between the hybridizing species across more granular taxonomic groups. We fit a mixed model with an interaction between taxonomic order and Jukes Cantor on the observed Patterson’s *D* value, with random effects of each study and species pair. The results, summarized in Figure 3, demonstrate that order specific slopes are supported for only a handful of taxa (significant taxon specific slopes shown with filled in labels), with increasingly weaker (smaller slope) relationships as more data is available per order. A similar approach for larger taxonomic units (classes and phyla) does not support phylum or class specific slopes. This analysis highlights two aspects. First, mammal orders tend to have positive slopes between genetic distance and introgression, potentially reflecting ancient introgression in mammals. Magnoliopsida (flowering plants) display a higher degree of heterogeneity, having the largest negative slope order (Ericales) and second highest positive slope (Brassicales). While taxonomic differences in introgression patterns are difficult to disentangle from noise due to low sampling and other systemic biases, we also note that within each taxonomic class, orders with more samples tend to show more negative slopes. More sampling is needed to further elucidate differences between taxa.

### Human associated organisms show elevated introgression

Human disturbance and association can increase the chances of both hybridization and introgression (van Hengstum, et al. 2012; Guo 2014; Ortego, et al. 2017; Grabenstein and Taylor 2018; Ottenburghs 2021). We annotated Patterson’s *D* values with human association status. A species was considered human associated if the source study mentioned that any of P1, P2, or P3 were hominids, domesticated species (or species in the process of domestication) or human pathogens, pests or parasites. We then fit a mixed model with human association as a fixed effect, and study and introgressing pair as random effects. Human association had a significant effect in this model, suggesting that species that are human associated had higher reported Patterson’s *D* values than those that are not. Prior work has suggested that hybridization is more common among human associated species (see Ottenburghs (2021) for a review), and so our findings fall in line with theoretical expectations. However, as Patterson’s *D* does not measure the direction or timing of introgression, it is hard to disentangle human mediated introgression through breeding domesticated species with wild relatives and introgression from domesticated taxa into native species. The latter may have strong negative consequences for native species and biodiversity (Todesco, et al. 2016), thus future studies should aim to identify the direction of introgression as well.

### Self-compatibility and sexual form influence introgression

Since introgression is inextricably linked to hybridization, drivers of hybridization are also likely to influence introgression. Previous work in plants has shown that a variety of life traits may influence the frequency of hybridization (Whitney, et al. 2010; Mitchell, et al. 2019). We therefore assessed whether life history traits within the largest dataset — flowering plants — had an effect on the magnitude of residual introgression. We annotated a subset of our flowering plant data with information about the self-compatibility (self-compatible, incompatible, or polymorphic) of each introgressing species pair as well as sexual form (dioecious vs hermaphroditic). Self-compatible plants are ones that are able to self-fertilize, and there may be differences in speciation rates between the two types of plants (Gervais, et al. 2011; Goldberg and Igić 2012; Roda and Hopkins 2019; Harkness and Brandvain 2020). In a mixed model including both self-compatibility of the introgressing species pair and sexual form as fixed effects, both show significant differences. Hermaphroditic plants show elevated introgression compared to dioecious (Supplementary Figure 7), while introgression is higher when one of the species is self-incompatible and the second is self-compatible compared to the rest (Supplementary Figure 8). Our usual caveats about potential reporting bias apply here as well. An important additional caveat is that these effects do not take into account phylogenetic non-independence of sexual forms and self-compatibility, and so could be driven by some other shared element of the different groups. Whitney, et al. (2010), for instance, find strong phylogenetic signal for the propensity to hybridize, an analysis we are unable to repeat here. However, our results present the first evidence that life history traits may impact introgression rates, motivating further study.

## DISCUSSION

One of the most enduring debates in evolutionary biology has been whether speciation can proceed with gene flow. To generally answer this question requires compiling data and timing of gene flow between species along a variety of taxa. More recent forms of the debate have taken the form of asserting that introgression might be a common feature of evolution (Seehausen 2004; Mallet, et al. 2016). Just as the formulation of the debate, we show that the field has changed in the approaches, both technical and statistical, that are used to detect introgression. Second, while we do not directly answer the question of the prevalence of gene flow in speciation, our meta-analysis demonstrates that introgression has left a mark on the genomes of extant populations and is supported by a vast number of studies across multiple eukaryotic systems. Finally, we find that there is extensive variation in the amount of remaining introgression across taxa. We discuss each of these considerations as follows.

### The evolution of the field

The study of introgression followed the same path as with many other subfields in the field of evolution, in which the underlying population genetic theory pre-dated the data resources necessary to test hypotheses. Despite speculation that hybridization provided raw materials for rapid adaptation, for example, such hypotheses were difficult to test without confirming that introgression was actually occurring (Anderson and Hubricht 1938; Heiser 1949; Anderson and Stebbins Jr 1954; Heiser 1973; Rieseberg and Wendel 1993). The sequencing revolution opened the floodgates for evolutionary biologists to interrogate the genome to answer specific questions about their system’s evolutionary history. The field as a whole has also advanced largely due to the capability to sequence the DNA of ancient individuals, as Patterson’s *D* statistic was initially formulated to detect gene flow between Neanderthals and contemporary humans, but was later fully derived to be extended for general use (Green, et al. 2010; Durand, et al. 2011). Our literature search identified a rapid increase in the number of papers with evidence for introgression over the last 15 years (Figure 1A). More importantly, the methods used to support introgression have shifted from purely qualitative approaches such as sequence similarity, to approaches that allow us to quantify the proportion of introgression such as *f*-statistics (Figure 1C). Introgression is also increasingly being found in taxa with no prior studies, but many Eukaryotic groups have had very little to no evidence of introgression (Figure 1B).

### Variation across taxa

Our data suggest several broad taxonomic patterns. Our mixed model fits show that there is more evidence for introgression in green plants than either animals or fungi. Within plants, this pattern seems to be driven by relatively few outliers from ferns and conifers (Polypodiopsida and Pinopsida), but less conservative models also indicate differences between fish (class Actinopteri) and many other taxa. The latter observation is consistent with previous work suggesting that fish have the highest rates of hybridization of any vertebrate taxa (Schwenk, et al. 2008). While further study is necessary, these differences could be driven by biology. There are good reasons for why taxa with faster rates of developing reproductive isolation, for instance mammals vs birds and anurans (Wilson, et al. 1974; Prager and Wilson 1975; Fitzpatrick 2004; Coughlan and Matute 2020; Matute and Cooper 2021), may show less evidence for introgression. Faster speciation leads to a smaller time-frame in which successful hybridization between divergent subspecies can occur. More rapid speciation also means that there are fewer fixed differences to introgress between species, and a higher amount of maintained ancestral polymorphism, increasing the ratio of ILS to introgression in statistics like Patterson’s *D*. On the other hand, rapid speciation may also be associated with increased introgression either due to introgression driving speciation, or rapid radiations leading to weak post-zygotic barriers in the resulting species complex (Mallet, et al. 2016). Speciation rates vary heavily both between phylogenetic classes and orders and within them (Rabosky 2009; Rabosky, et al. 2013; Schluter and Pennell 2017; Coughlan and Matute 2020). Furthermore, there is a great degree of variation in the amount of sympatry (Nosil 2013; Matute and Cooper 2021) and overall hybridization rates (Chen, et al. 2018; Mitchell, et al. 2019). Thus, we expect to see variation in Patterson’s *D* across eukaryotes, at varying scales, due to a variety of biological phenomena.

By linking observed Patterson’s *D* values with genetic distances calculated from publicly available data, we were able to ask several questions about introgression and divergence. First, we find a negative relationship between genetic distance and Patterson’s *D* (Figure 2B). While this result is largely expected from several theoretical perspectives (Hamlin, et al. 2020), it is overall a weak relationship. As species diverge, build-up of reproductive isolation presents both fewer opportunities for introgression (Harrison and Larson 2014; Kenney and Sweigart 2016) and increases the selection against introgressed regions (Staubach, et al. 2012; Jagoda, et al. 2018; Petr, et al. 2019). However, several biases in Patterson’s *D* and the study of introgression may explain the relatively weak relationship, and we discuss these further in the Caveats section of this discussion. However, this overall weak pattern may be driven by looking for a single slope of introgression vs genetic distance across all studied taxa, with different slopes canceling each other out.

**Figure 2:**
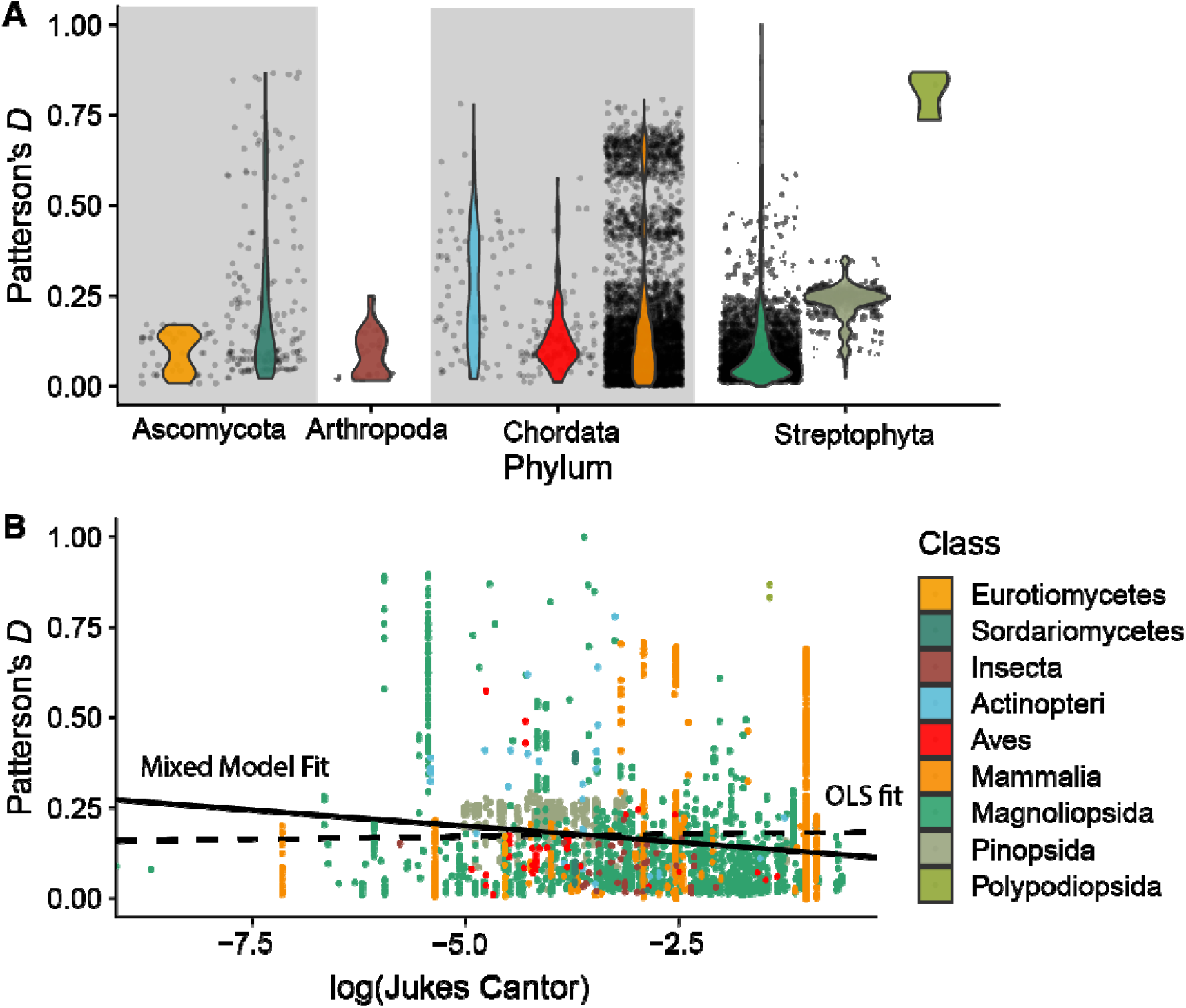
A) Distribution of Patterson’s *D* values across phyla and classes. B) Weak negative relationship between genetic distance and Patterson’s *D*. Solid line – mixed model fit with study and species pair as random effects. Dashed line – ordinary least squares fit.

To examine the latter case, we calculated the slope of the best fit linear models between genetic distance and Patterson’s *D* across individual taxonomic orders (Figure 3). Unsurprisingly, the amount of observations in an individual taxon played a strong role in determining the slope of the relationship between genetic distance and Patterson’s *D*. As more data becomes available for any order, the slope becomes less steep, but many orders show positive, rather than negative, relationships between Patterson’s *D* and genetic distance. Primates, for example, have a fairly strong signal for increased divergence leading to increased evidence for introgression, but this is likely biased by the heavy focus on ancient introgression in hominids, giving many positive values of Patterson’s *D* for relatively highly diverged species pairs. Similarly, increased signals of introgression between diverged taxa can be a result of human association, and mixed model fits do support significantly higher introgression reported among human associated lineages (Supplementary Figure 6). Introgression is often artificially generated between diverged species in agricultural settings, and interest in ancient introgression may be higher for domesticated taxa, leading to human associated taxa showing different patterns compared to other species. Two major lines of inquiry are suggested by our data. First, the effects of ancient introgression in determining the relationship of Patterson’s *D* and genetic distance need to be explored. While the general expectation has been for an overall decrease in introgression as taxa diverge (Roux, et al. 2016; Hamlin, et al. 2020), it is possible that this signal is swamped by ancient introgression, or that for some taxa introgression is more likely between diverged species pairs. Second, it seems that there may be genuine differences in the relationship between genetic distance and introgression among some of the best studied taxa (Figure 3), a pattern that, to our knowledge, has not been previously reported or expected. These differences may be driven by study effort differences, but also due to evolutionary history of introgression or differences in the process of speciation between taxa.

**Figure 3:**
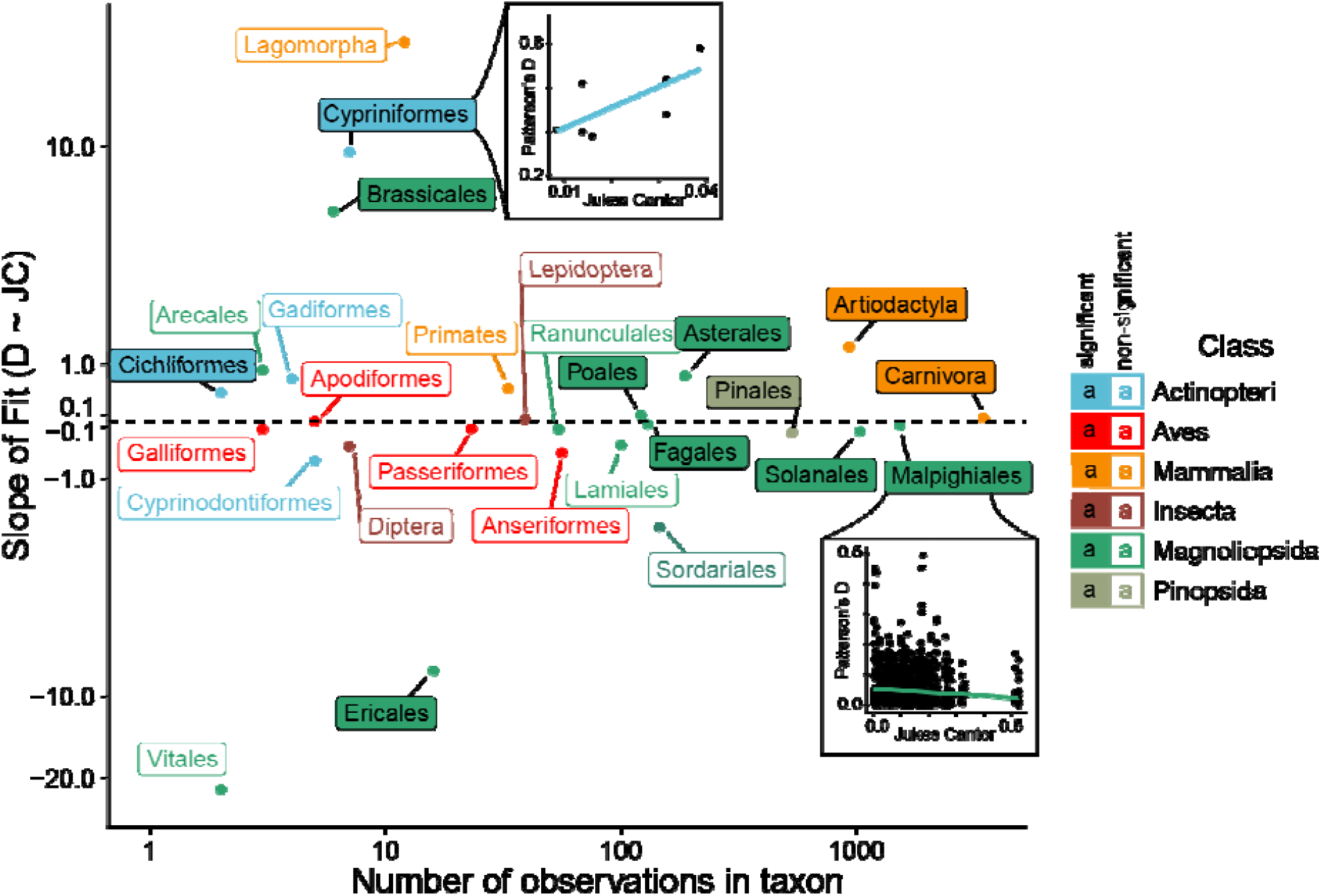
Increasing observations per taxon suggest weaker relationships between genetic distance and Patterson’s *D*. Results from a mixed model fit with interactions between genetic distance (measured as Jukes Cantor) and taxonomic order. Significant interactions of order and slope of Jukes Cantor are represented by filled in boxes, non-significant with empty. Orders with two or fewer representative species pairs are excluded.

### Recent vs ancient introgression

Our dataset is unable to distinguish between ongoing/recent and ancient introgression. Even when Patterson’s *D* is applied correctly, it can detect both ancient and recent introgression. Ancient introgression in a taxon may lead to many species pairs with positive Patterson’s *D* values reported in our data-set (see Pines, for instance), while a recent introgression event is likely to be represented by just a single species pair. This generates a potential bias for elevated Patterson’s *D* between more diverged populations, as Patterson’s *D* identifies signals of introgression between pairs of species rather than at a particular branch/timepoint. Our lack of ability to time introgression in this study also limits us from knowing whether we are seeing introgression between mostly reproductively isolated species, or just the signals of ongoing speciation with gene flow. The solution for comparative biology of introgression is to identify not only the proportion of introgression, but the timing as well. Several methods are making progress on this front (Edelman, et al. 2019; Shchur, et al. 2020; Martin and Amos 2021; Svedberg, et al. 2021), while some *f*-statistics approaches can identify likely introgression timing given a tree topology (Malinsky, et al. 2018). The future of the field thus may be better able to deal with some of the caveats we discuss next.

### Caveats

Our results are not devoid of caveats. One of the main findings of our analyses is the extensive variation in the depth and quality of reports claiming support for introgression. While the field has uniformly moved towards the study of introgression using genome sequences, not all studies have used whole genome analyses. This is due to the extreme genome size of some taxa (Gregory 2021) and because there are potential trade-offs in the number of individuals sequenced and the amount of genome sequenced. The benefits and caveats of reduced sequencing has been described elsewhere (Puritz, et al. 2014; Lowry, et al. 2017), but briefly, the selection of markers will invariably bias estimates of introgression as reduced representation sequencing (RRS) data inherently underestimates true levels of diversity (Gautier, et al. 2013; Cariou, et al. 2016). Moreover, comparing the metrics of introgression revealed by these different methodologies has proven challenging as our linear mixed models showed that incorporating this variable as a random effect explains a substantial portion of the model variance.

A second noteworthy caveat pertains to the limitations of Patterson’s *D*, the statistic that is most widely available. First, because of its very proposition Patterson’s *D* detects the excess of introgression into one species. If the donor species is contributing the same alleles to two sister species, then Patterson’s *D* will be zero. On the other hand, the metric is not reliable when one of the taxa has experienced a bottleneck, and it produces false positives under certain demographic scenarios (Martin, et al. 2015; Hibbins and Hahn 2021). Finally, the statistic relies on a specific species tree being true — when the test is applied to populations that may not meet the topology expectation, it is likely to return meaningless values. Since species relationships in taxa with high a degree of introgression are hard to determine (root node of Neoaves, for example (Prum, et al. 2015)), such errors might ironically be more prevalent for taxa in which introgression actually has occurred.

These caveats extend to our analysis of the relationship between genetic distance and Patterson’s *D*. First, Patterson’s *D* is not suited to measuring the proportion of introgression, which is expected to decrease with increasing genetic distance. Instead, it is a measure of the presence of introgression, and ancient introgression may therefore generate a pattern of relatively flat rates of introgression across genetic distances. Second, our dataset consists of a variety of introgression events, some recent or ongoing, others quite ancient. On the other hand, our data is also depleted for small/zero values of introgression, as researchers are unlikely to measure introgression between distantly related taxa that are unexpected to have a history of hybridization, but also because researchers are unlikely to report small values of Patterson’s *D* due to the “drawer effect” (Scargle 1999).

Perhaps the largest difficulty does not pertain to Patterson’s *D* itself but to how the results of the tests are reported. The lack of consistency in reporting introgression paints a muddled picture of its frequency and any differences between taxa. In terms of reporting, a variety of approaches are used — in some cases, researchers report only those values of introgression statistics that represent the particular set of introgression events under study. In others, only significant values of statistics are reported (which naturally leads to a depletion of low values of Patterson’s *D*). Sometimes a mix of approaches is used, where only some particular sets of species are tested for introgression and only some values are reported. We argue that reporting all possible Patterson’s *D* values given the groups under study would facilitate future comparative studies. We strongly suspect that the banding observed in Figure 2A and others is caused by the subjective application of significance thresholds. This issue is related to the potential of a ‘drawer effect’, with differing values across papers being considered significant enough to report. Lastly, it is nearly impossible to disentangle differences in reporting and study effort between fields from actual differences in introgression frequency. While studies such as this one are helpful to identify general trends, until the field unites behind a unified reporting standard, truly comparative studies that use a single set of approaches to interrogate introgression across taxa will be necessary.

### Directions for the field

Alongside the development of our understanding that speciation is a process and not an event has come the appreciation of ongoing gene flow between what are often believed to be good species. Introgression, rather than an exceptional occurrence, seems to be a common feature of evolution in Eukaryotes, at least in cases where it has been sought. Studies across a wide array of eukaryotes are now shedding light on the frequency of introgression. However, several developments are necessary to understand the drivers of introgression.

#### A unified reporting standard

In order to answer the overarching question of “How prevalent is introgression across the tree of life?” researchers must either shift their focus from taxa-centric studies of introgression to more clade-centric studies (e.g. Malinsky, et al. (2018); Edelman, et al. (2019); Hamlin, et al. (2020); Small, et al. (2020); Suvorov, et al. (2021)), however, we recognize that this may not always be possible or feasible. Alternatively, we suggest the following unified reporting standard, to further advance the field’s abilities to perform comparative analyses of introgression across the tree of life. We first suggest that researchers report a genome-wide Patterson’s *D* value, number of ABBA sites, and number of BABA sites for all possible pairwise comparisons of groups that don’t violate the assumed species tree topology. Although Patterson’s *D* has its shortcomings it is very simple to compute (see ANGSD (Korneliussen, et al. 2014), scikit-allele (Miles 2020), or D-suite (Malinsky, et al. 2021)) and can be calculated from population genetic data as well as whole genome alignment data, which makes it applicable to test for the presence of introgression on both population level and phylogenetic time scales. Additionally, we recommend researchers assess significance using a standard block jackknife procedure — as first described in Reich, et al. (2009) — and subsequently report the standard error and corresponding Z-score. Secondly, we suggest that researchers calculate a genome wide 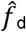 value (Martin, et al. 2015) for only statistically significant Patterson’s *D* configurations, since, in the absence of the known demographic history of one’s study system, there is no formal way to assess significance for the inferred admixture proportion; subsequently an 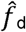 value for an insignificant Patterson’s D configuration is uninterpretable. Genome-wide 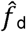 has been shown to be a robust and conservative estimator of the genome wide admixture proportion (Martin, et al. 2015; Pfeifer and Kapan 2019; Pfeifer, et al. 2020). Furthermore, as an analog of Patterson’s *D* it can also be applied to both population genetic data as well as whole genome alignment data. We would also like to emphasize that this unified reporting standard should not replace any new methods to detect and/or quantify introgression, but instead provide the minimum and necessary information to empower future comparative studies. Indeed, new methods to quantify the timing of introgression are likely to increase the power of comparative studies by identifying the timing and direction of introgression.

## Conclusions

Our goal with this piece is not to become the last word on the question of the prevalence of introgression across taxa in nature. Instead, we provide a state-of-the-art compilation that reveals the current understanding of the field, tests current hypotheses and most importantly highlights the most notorious gaps in the field. We find that introgression has been identified across Eukaryotes, but sampling is uneven, and reporting needs to be standardized to allow for comparative questions in introgression to be answered. Although our dataset is not able to answer these questions, we find several patterns that motivate further study. The last 15 years have seen a transformation in the way we study introgression, the next 15 will hopefully reveal much about its drivers and prevalence.

## Supporting information

Supplementary Tables 2-9, Files 1-2 and Copies of Supplementary Figures 1-9

## ACKNOWLEDGEMENTS

We would like to thank Yaniv Brandvain, Senay Yitbarek, and Adam Stuckert for useful conversations and advice. This work was supported by by the National Institute of General Medical Sciences of the National Institutes of Health (NIH) under Award Number R01GM121750 to DRM and DP was supported under Award Number 1R35GM128946-01.

## DATA AVAILABILITY

All scripts/data used for analyses and to generate plots are available on request, and will be made available on Dryad prior to publication.

## METHODS

### Search criteria

To identify the taxa in which introgression has been studied, and create a comprehensive list of papers from which we could extract Patterson’s *D* values, we performed a Web Of Science search. We first searched for papers which contained the terms “introgression”, “hybrid” and “genomic”, and complimented the results with any papers citing any of the several papers that defined major *f*-statistics (Green, et al. 2010; Martin, et al. 2015). Due to the relative breadth of our initial search criteria, we captured many papers on experimental introgression lines, hybrids occurring solely in the lab, methods to detect introgression or hybridization and many perspectives and reviews. Papers were then manually inspected for claims of introgression, resulting in nearly 724 papers with claims of introgression. These papers were annotated for the major taxonomic group of the study organism as well as the types of evidence provided when introgression was confirmed. The list of these contributions appears in Supplementary File 1.

### Introgression test classification

Since a large variety of methods are used to detect introgression, we binned them into six categories depending on the type of information they use. The first category, sequence similarity, consisted of direct sequence comparisons, whether by genetic distance, *F*_ST_, *d*_xy_, or sequence alignments. The “cline” group consisted of any studies that used clinal data to identify loci that introgress past the contact zone. We included both studies that used geographic and genomic clines. The tree group included any studies in which gene trees were used to identify introgression. This could be mitochondrial-nuclear mismatch, differences in trees across several nuclear genes or use of software like TreeMix (Pickrell and Pritchard 2012) or Twisst (Martin and Van Belleghem 2017) or QuIBL (Edelman, et al. 2019). The clustering group consists of methods such as NEWHYBRIDS (Anderson and Thompson 2002), STRUCTURE (Pritchard, et al. 2000), fastSTRUCTURE(Raj, et al. 2014), ADMIXTURE (Alexander, et al. 2009; Alexander and Lange 2011) or simply PCAs of genetic diversity to identify hybrid and admixed individuals. Studies which included some form of demographic model fitting to identify ongoing admixture were included in the demography group. Finally, any studies that included the many types of *f*-statistics listed in Table 1 were included in the *f*-statistics group.

### Extracting *f*-statistics and criteria for inclusion

Since our goal was to quantify the strength of evidence for introgression in different taxa, we next examined any papers with some form of *f*-statistics to extract data. We only included genome-wide Patterson’s *D* values. For each study, we extracted the populations under study, their reported *f*-statistic and its value and reported significance. Due to high variability in which statistics were reported, we also annotated the genomic data type (whole genome sequencing, RAD/GBS, transcriptome/exome, amplicon sequencing) used for the study as well as whether the authors reported all possible *f*-statistics, only significant ones, or a specific subset of interest, as well as whether multiple outgroups were used. We first pruned the data for significance. Patterson’s *D* values above 0.05 had to be significant at the p < 0.05 or Z > 3 level. The power of Patterson’s *D* relies on the number of sites showing either an ABBA or BABA configuration. To obtain significant, but very small, Patterson’s *D* values, a study therefore has to identify very large numbers (but small differences) in the number of ABBAs and BABAs. As a result, most studies are under-powered to detect extremely low levels of introgression; we included all values of Patterson’s *D* < 0.05 in the remainder of the analyses.

We next filtered our data for Patterson’s *D* values that are likely to represent the true species topology. When Patterson’s *D* statistic is negative, introgression is supported between populations 1 and 3, while positive values indicate introgression between populations 2 and 3 (Supplementary Figure 1). To allow easier comparisons between studies we first re-ordered all species triplets so that the *D*-value measured introgression between populations 2 and 3 only, giving us only positive Patterson’s *D* values. We next reorganized the dataset to use only those configurations of populations 1, 2 and 3 that displayed the weakest evidence of introgression. When the wrong species tree topology is used in calculating Patterson’s *D*, it may result in highly elevated values, detecting introgression where shared ancestry is responsible (Supplementary Figure 1). We reasoned that when multiple tree topologies were considered in our dataset, the smallest *D*-statistic would correspond to the most likely “true” species relationship, and present the most conservative view of introgression. These filters reduced our dataset to 18,339 values of Patterson’s *D* across 112 studies, although including all observations did not qualitatively change our results (data not presented).

### Mixed Model Selection

To account for the random effects stemming from differences in reporting (all possible pairs, only significant, or a specific subset, Supplementary Figure 5) and power of different genomic sequencing (Supplementary Figure 4), we used several mixed modeling approaches. In our most conservative analysis, we include the source study for each value as a random effect. In this approach, the random effect of study accounts for nearly half of the residual variation in observed Patterson’s *D* values. However, as each study was generally limited to an individual taxon, this conservative approach is likely to under-power our ability to detect meaningful differences between biological groups. As a second approach, we included random effects for the sequencing type used in each study as well as a second random effect for the reporting type. Sequencing type accounts for roughly 17% of variance, while reporting type accounted for just over 5% of observed variability in Patterson’s *D*. Lastly, to account for phylogenetic non-independence of observed introgression statistics, we included a random effect of the species pair in both approaches, coupled with a fixed effect of genetic distance (calculations explained below). We fit all models using the lme4 package in R 4.0.3 (Bates, et al. 2015; R Core Team 2020), while pairwise comparisons between fixed effects were performed using the emmeans package (Lenth 2020).

#### Testing the effect of genetic distance on residual introgression

Since introgression may be impacted by genetic distance between species pairs, we attempted to obtain measures of divergence for all species pairs with measures of Patterson’s *D*. We first identified the NCBI taxon id for each P2 and P3 using a custom script and NCBI’s e-utilities tools. The breadth of our data meant that no individual gene could be used to measure divergence between the majority of our species pairs. As a result, we downloaded up to 10,000 sequences from NCBIs nucleotide database (2018) for each species in the pair. Reciprocal best BLAST hits (Camacho, et al. 2009) from the two species’ sequences were then aligned using CLUSTAL (Sievers and Higgins 2018), and average Jukes-Cantor distance was calculated for the resulting alignments in R 4.0.1 using the ape package (R Core Paradis and Schliep 2019; Team 2020). 9,804 of our Patterson’s *D* values were successfully annotated with genetic distance in the process. The resulting genetic distances were then used as a fixed effect in our mixed modeling approaches.

#### Effect of human association on introgression

We annotated each Patterson’s *D* entry with human association status manually. The source studies of each Patterson’s *D* value were examined for statements whether any of the species in the study are domesticated, in the process of domestication, human pathogens, pests or parasites, or simply hominids. We first included human association in models 1-30 (Table 2), but it was not a significant fixed effect in any of them. This is because there is a high degree of overlap between phylogenetic order and human association. Most of the Primate studies are on hominids, while nearly all Artiodactyla studies regarded domesticated animals. Since we are unable to separate out taxonomy and human association we instead fit human association by itself. We fit a mixed model with human association as a fixed effect, and genomic data type, reporting type and introgressing pair identity as random effects. We find that human association is a significant effect in this model in a post-hoc test following ANOVA. Models were again fit using lme4 in R version 4.0.1.

#### Effect of life history on signal of introgression

Finally, we studied whether life history components had an effect on the amount of residual introgression. We focused on flowering plants because they have broad sampling across many taxa and there are pre-existing expectations about introgression and the evolution of self-compatibility (Harkness and Brandvain 2020). We downloaded data on the self-compatibility and sexual form of plants from the Tree of Sex database (Bachtrog, et al. 2014). Species with records in the Tree of Sex database were then annotated in our dataset to include self-compatibility status, sexual system and annual/perennial status. We then examined whether self-compatibility influenced introgression by categorizing each introgression event as between two self-compatible species, one self-compatible and one not, or both self-incompatible (Supplementary Figure 8). We fit a mixed model with the pair’s self-compatibility status and sex system (dioecy vs hermaprhoditism) as fixed effects without interactions, and study and pair as random effects. Models were fit in R using lme4, and significance of comparisons between levels of fixed effects was determined using least-squares means calculated using the emmeans package.

## SUPPLEMENTARY MATERIAL

**Supplementary Figure 1:**
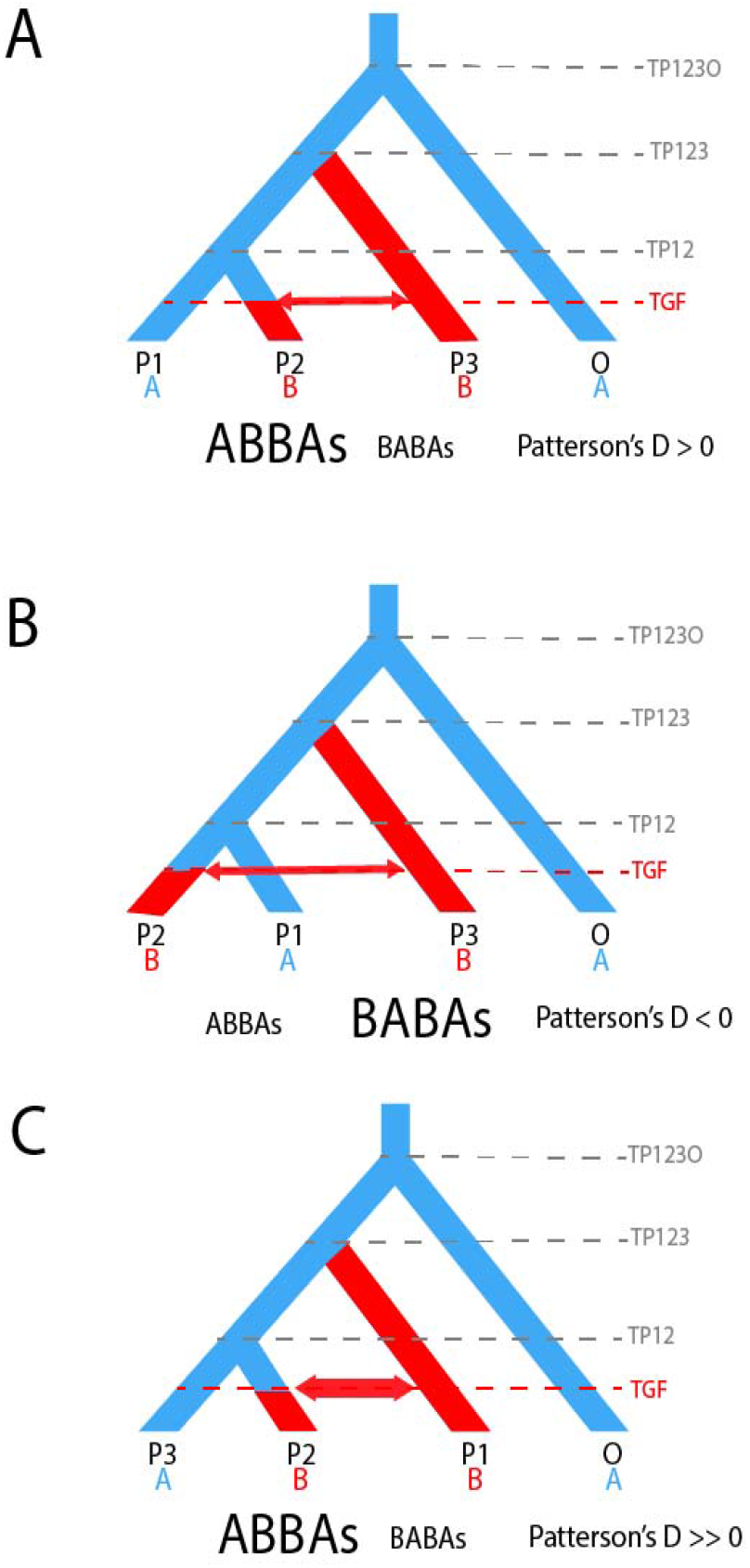
Patterson’s *D* expected under various scenarios. In all panels lineages in blue carry the ancestral allele and are represented by an A, lineages in red carry the derived allele and are represented by a B, TP123O represents the time when the outgroup population diverged from the ancestral population P123, TP123 represents the time when P3 diverged from the ancestral population P12, TP12 represents the time when P1 and P2 diverged, and TGF represents the time when gene flow occurred. A) The true species relationship is shown, with sister taxa P1 and P2, and introgression between P3 and P2. Introgression increases the shared derived alleles between the two populations (ABBAs), leading to an overall positive Patterson’s D. B) If P2 and P1 are swapped, Patterson’s D becomes negative, so changing the sign of negative Patterson’s D and swapping the identity of P1 and P2 allows for easier comparisons. C) When the wrong tree topology is used, Patterson’s D values can be highly misleading. In this case, P1 and P2, which are actually sister taxa, will share many derived alleles, leading to a highly elevated Patterson’s *D*. When multiple arrangements of a triplet are available, we assume the arrangement with the lowest Patterson’s *D* is most likely to be the “true” species tree.

**Supplementary Figure 2:**
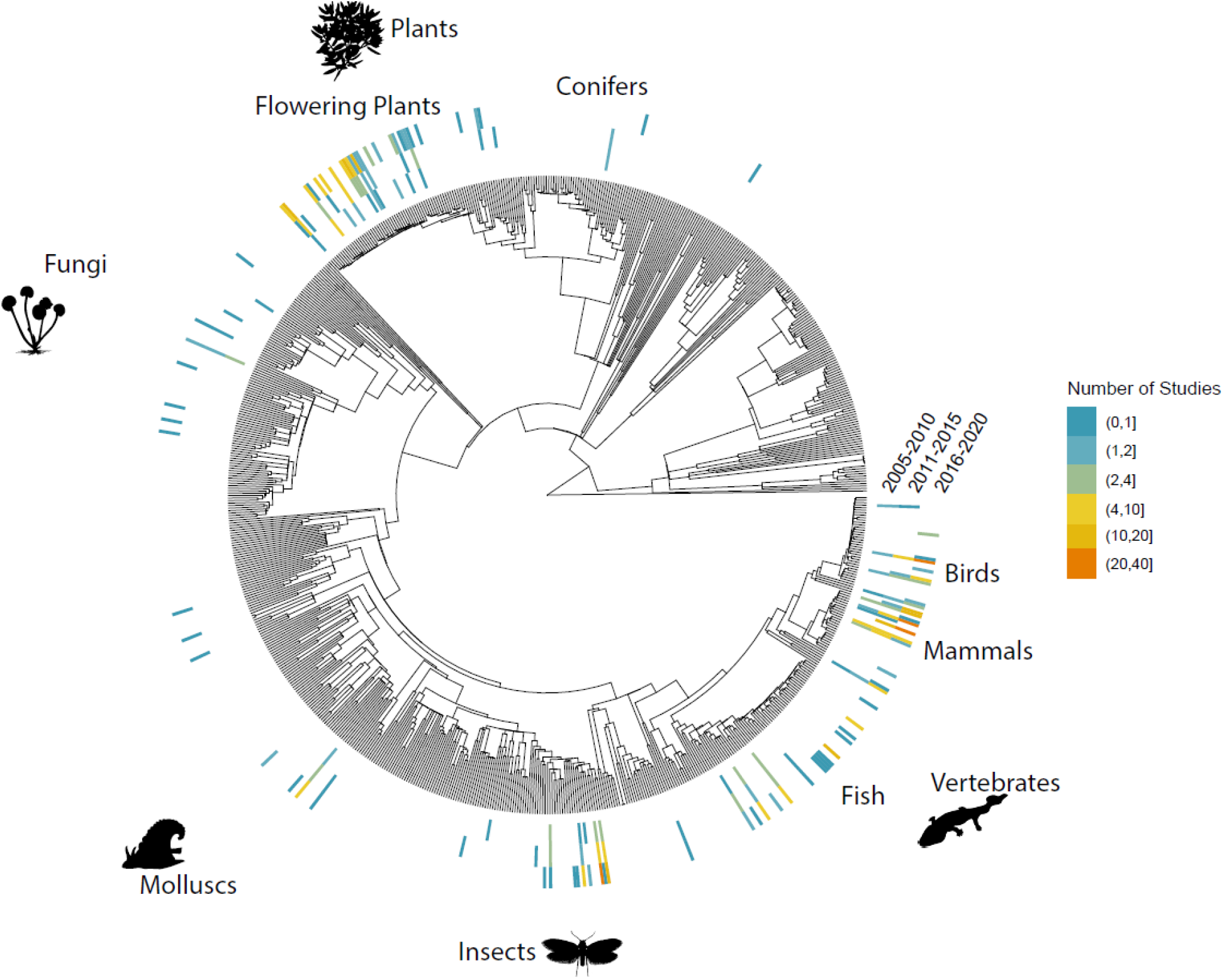
Number of introgression studies across eukaryotic orders in 5 year increments since 2005. Overall, the number of systems with evidence for introgression has vastly increased in the years 2016-2021 (outer ring), with many orders that had no prior evidence of introgression seeing publications (inner two rings).

**Supplementary Figure 3:**
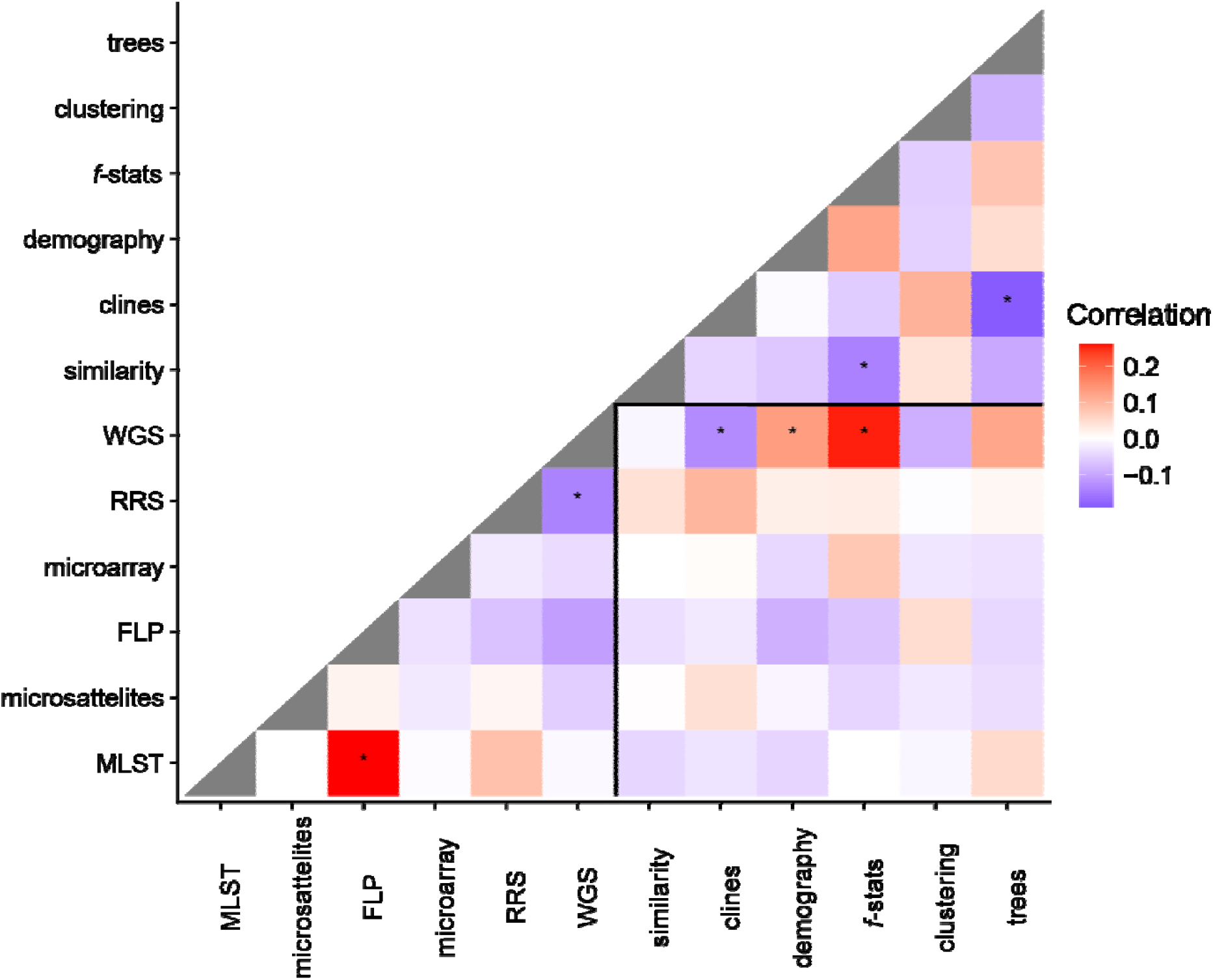
Correlations between various data types and evidence for introgression. Papers could include both multiple data types and types of evidence. Significance of correlation indicated by asterisk (p<0.05), following Bonferroni correction.

**Supplementary Figure 4:**
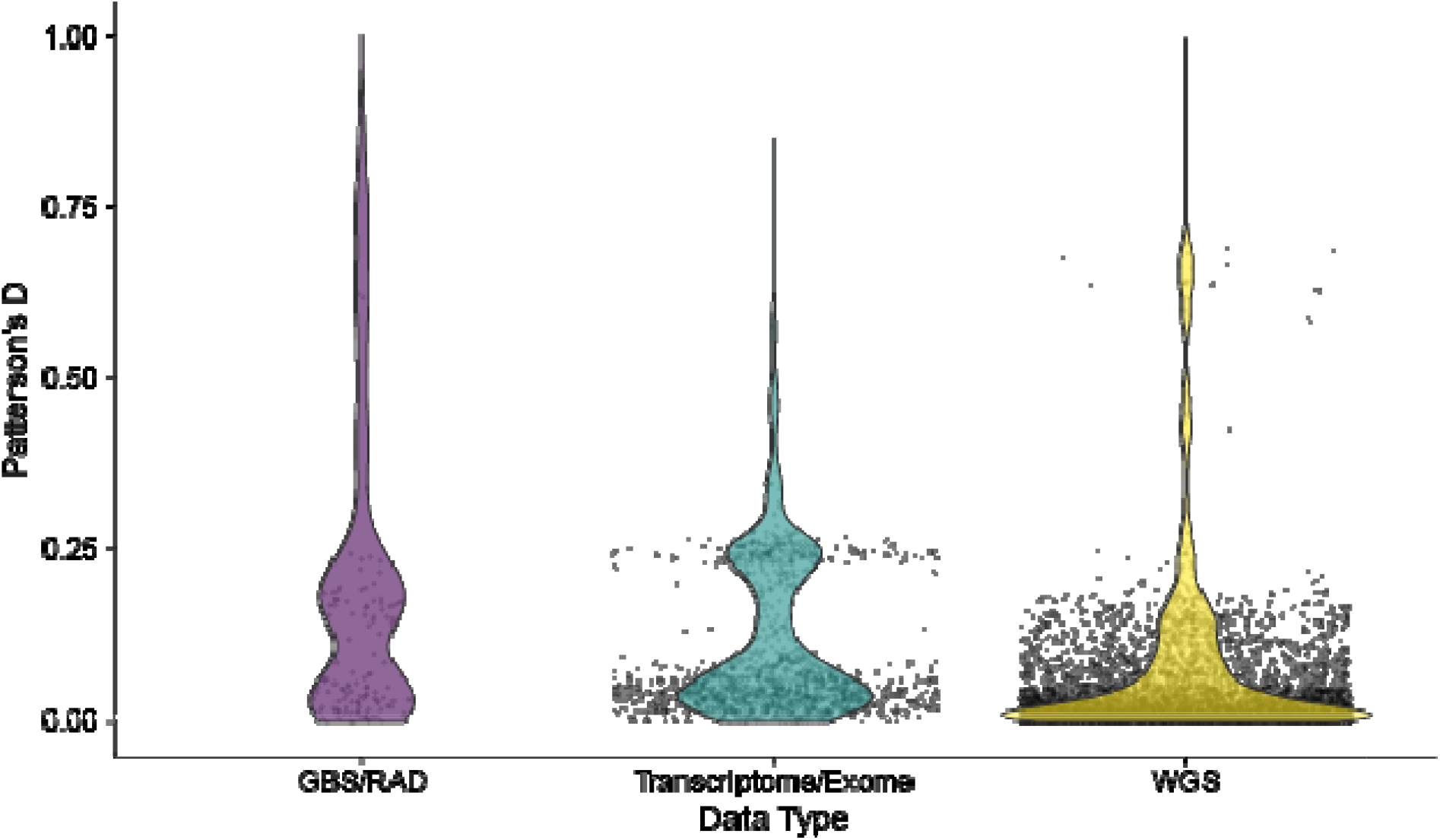
Distribution of Patterson’s *D* vs sequencing type: We excluded a few data points from amplicon or mixed data. While a simple t-test shows significant differences between data types and reported Patterson’s D, the difference is not significant in a mixed model accounting for the random effects of each study. Since no single study reported values from multiple data types, it is impossible to disentangle whether different sequencing methods have different error rates in detecting introgression. Banding may be resulting from differences in reporting criteria and significance levels of different papers.

**Supplementary Figure 5:**
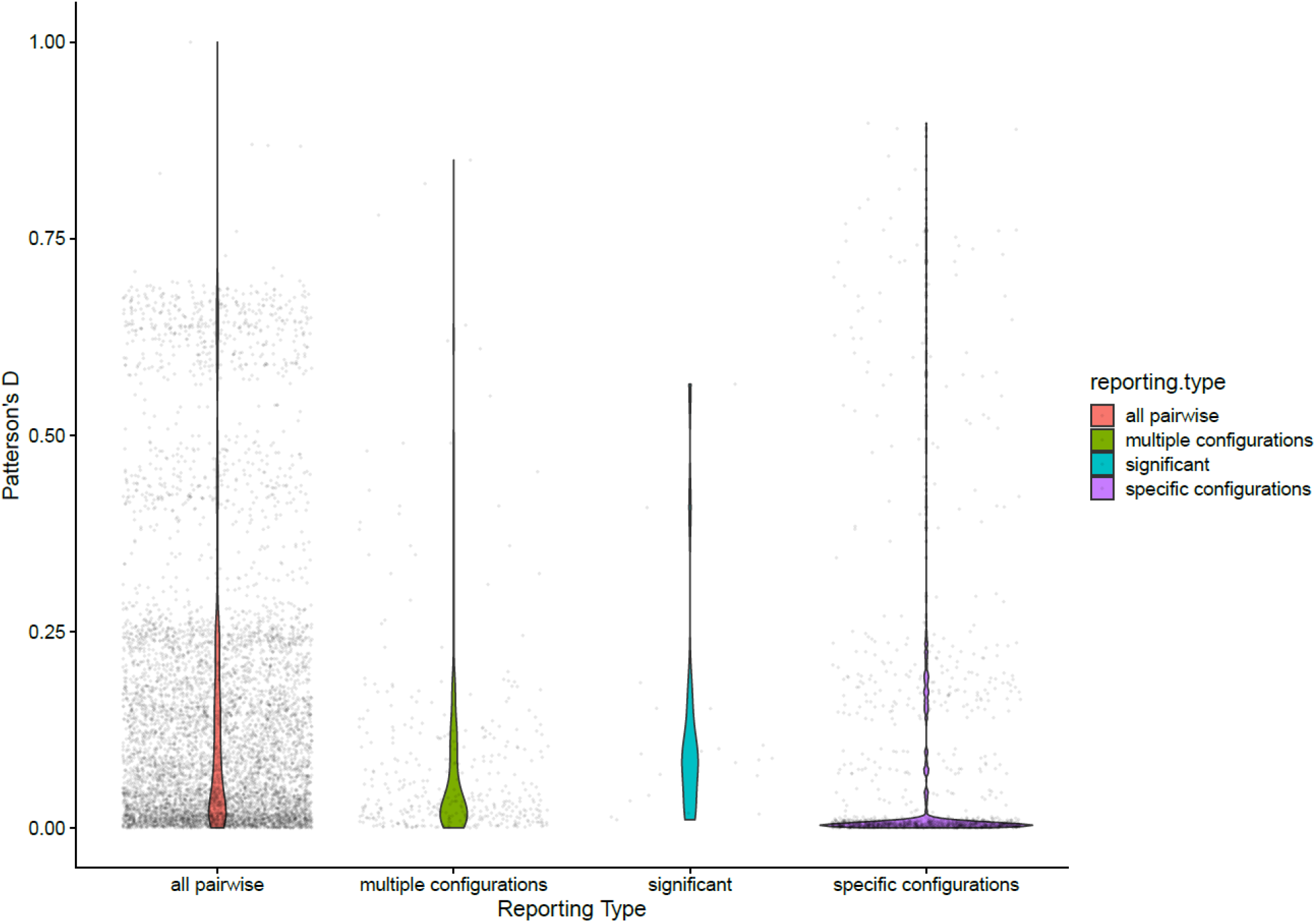
Distribution of reporting types. Papers were classified based on whether they reported all pairwise reports between studied taxa, multiple different configurations, only significant configurations, or specific configurations of interest.

**Supplementary Figure 6:**
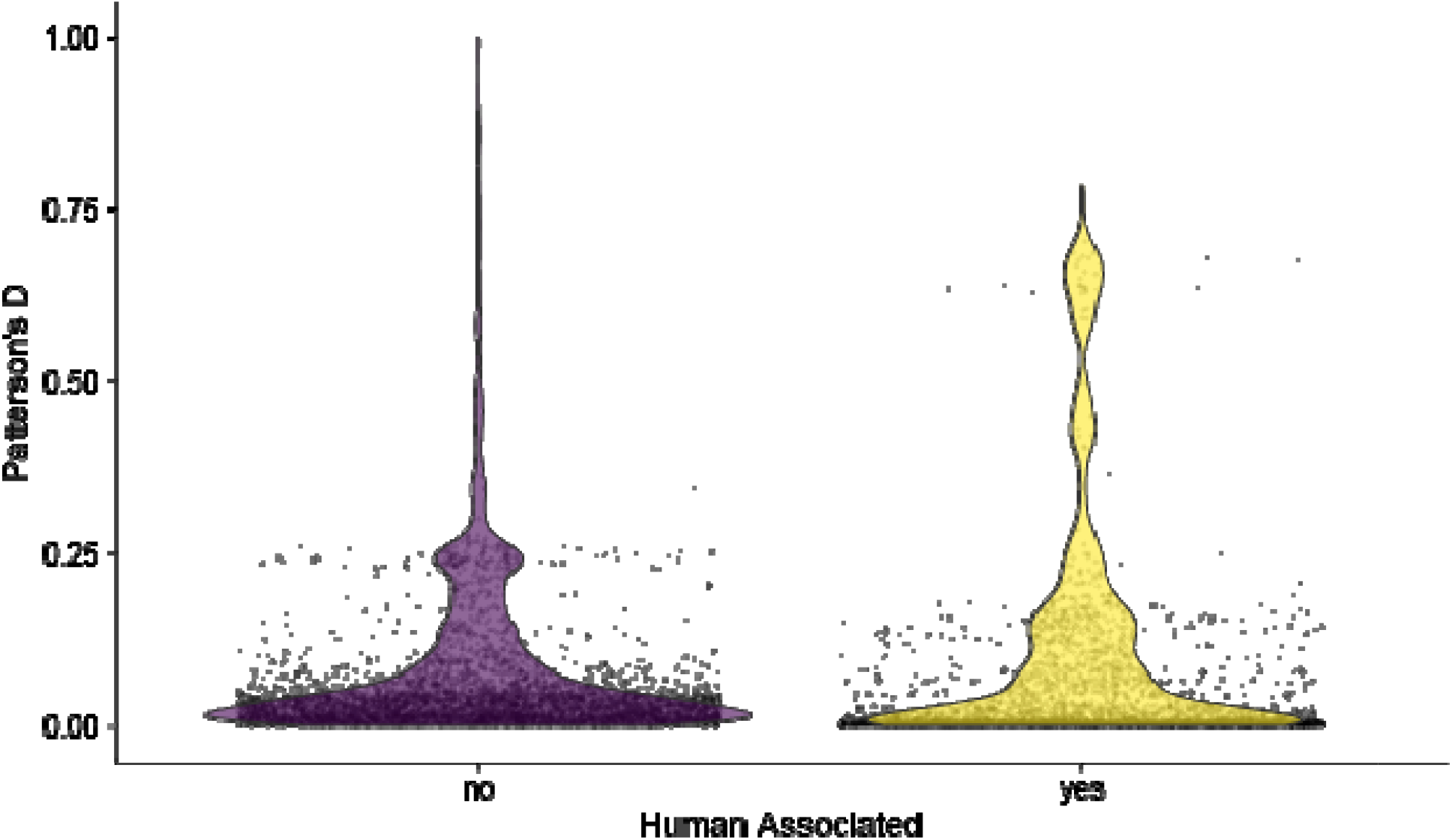
Distribution of Patterson’s D vs Human association status: Taxa that are associated with humans (agricultural animals/plants, pests and diseases) show elevated Patterson’s D values compared to taxa without clear human association in a simple t-test. However, these differences are insignificant once random effect of study is included, because no studies look at introgression in both wild and domesticated systems at the same time.

**Supplementary Figure 7:**
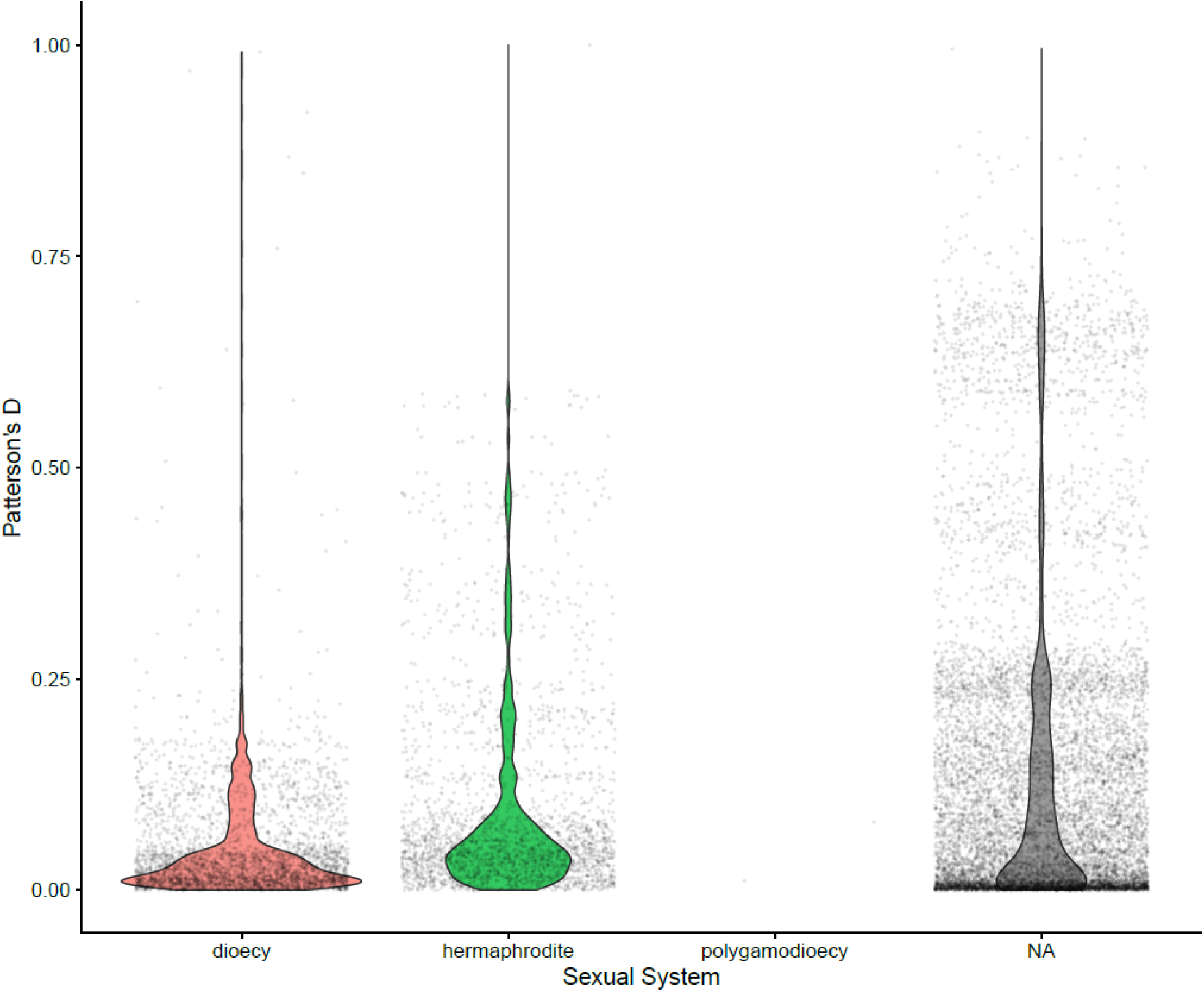
Sexual systems and Patterson’s *D*. Elevated Patterson’s D is seen in hermaphroditic taxa compared to dioecious among our flowering plant data set. All introgressing species pairs shared sexual system except for two samples of polygamodioecious species. Sexual system annotations were not available for a large proportion of the data.

**Supplementary Figure 8:**
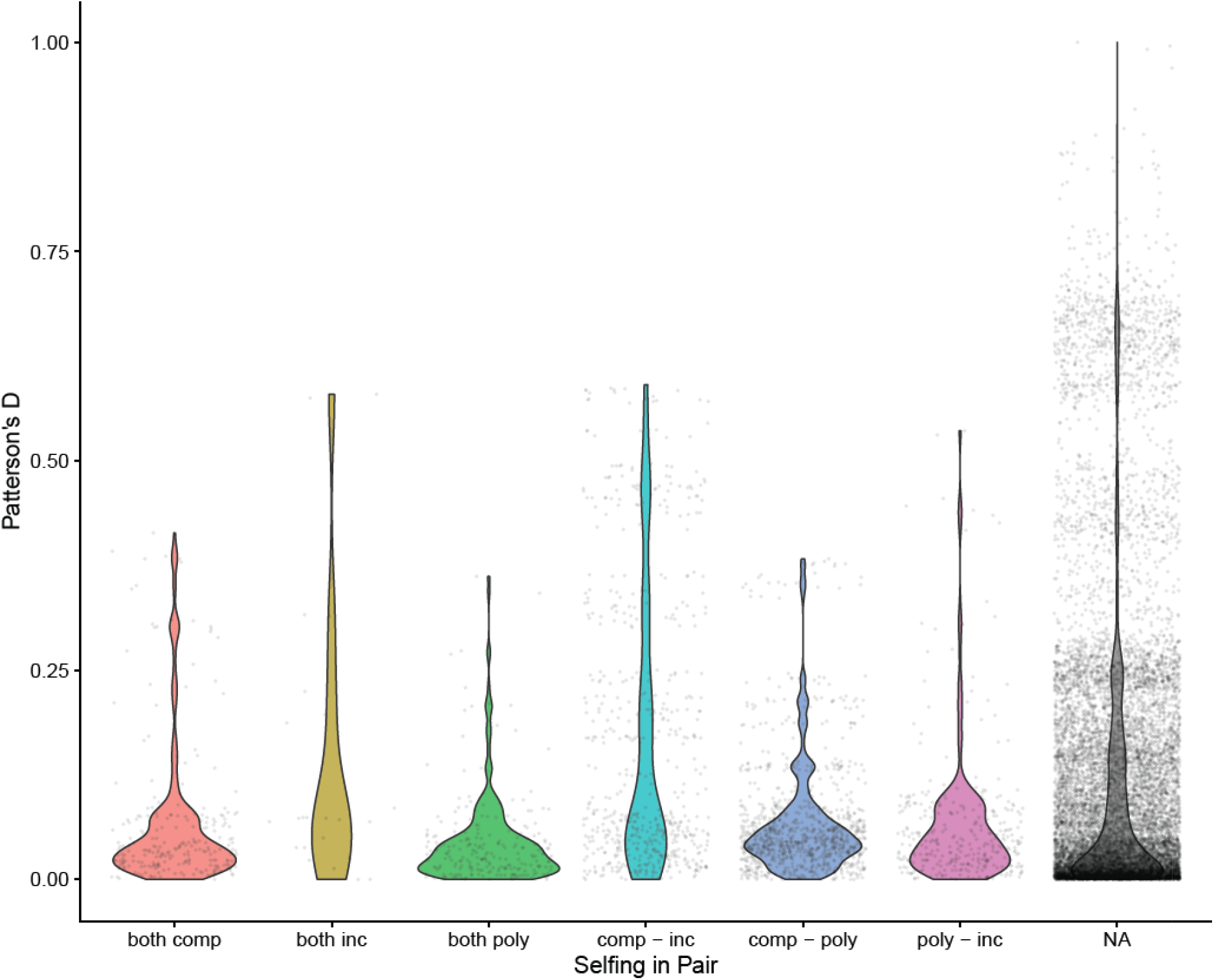
Patterson’s *D* values over various configurations of selfing among flowering plants. Comp=self-compatible, poly=polymorphic in species, inc=self-incompatible. Mixed model fits show significantly increased Patterson’s D for comp-inc class only. Self-compatibility annotations were not available for a large proportion of the data.

**Supplementary Table 1:** Number of papers with evidence for introgression by taxonomic order, top 30 orders.

| **ORDER** | **NUMBER**<br>**OF PAPERS** |
| --- | --- |
| PRIMATES | 51 |
| PASSERIFORMES | 43 |
| POALES | 37 |
| ARTIODACTYLA | 35 |
| DIPTERA | 31 |
| LEPIDOPTERA | 29 |
| RODENTIA | 22 |
| ANURA | 21 |
| LAMIALES | 19 |
| ASTERALES | 18 |
| CARNIVORA | 17 |
| MALPIGHIALES | 17 |
| FAGALES | 16 |
| BRASSICALES | 15 |
| PINALES | 15 |
| SQUAMATA | 15 |
| CICHLIFORMES | 14 |
| MYTILIDA | 14 |
| CYPRINIFORMES | 13 |
| HYMENOPTERA | 12 |
| ROSALES | 12 |
| SALMONIFORMES | 12 |
| LAGOMORPHA | 11 |
| URODELA | 11 |
| FABALES | 10 |
| GALLIFORMES | 9 |
| ANSERIFORMES | 8 |
| SACCHAROMYCETALES | 8 |
| SOLANALES | 8 |
| ASPARAGALES | 7 |

Supplementary Tables 2-9: Available in a separate file

## References

2018. Nucleotide [Internet]. Database resources of the National Center for Biotechnology Information. Nucleic Acids Research 46:D8–d13.

Alexander DH, Lange K. 2011. Enhancements to the ADMIXTURE algorithm for individual ancestry estimation. BMC Bioinformatics 12:246.

Alexander DH, Novembre J, Lange K. 2009. Fast model-based estimation of ancestry in unrelated individuals. Genome Research 19:1655–1664.

Anderson E, Hubricht L. 1938. Hybridization in Tradescantia. III. The Evidence for Introgressive Hybridization. American Journal of Botany 25:396–402.

Anderson E, Stebbins Jr GL. 1954. Hybridization as an evolutionary stimulus. Evolution:378–388.

Anderson E, Thompson E. 2002. A model-based method for identifying species hybrids using multilocus genetic data. Genetics 160:1217–1229.

Arnold ML, Martin NH. 2009. Adaptation by introgression. Journal of Biology 8:82.

Avise JC. 2004. Molecular markers, natural history and evolution, 2nd edition: Sinauer Associates.

Bachtrog D, Mank JE, Peichel CL, Kirkpatrick M, Otto SP, Ashman TL, Hahn MW, Kitano J, Mayrose I, Ming R, et al. 2014. Sex determination: why so many ways of doing it? PLoS Biology 12:e1001899.

Barton NH. 2001. The role of hybridization in evolution. Molecular Ecology 10:551–568.

Barton NH, Gale KS. 1993. Genetic analysis of hybrid zones. In: Harrison RG, editor. Hybrid zones and the evolutionary process. p. 13–45.

Barton NH, Hewitt GM. 1989. Adaptation, speciation and hybrid zones. Nature 341:497–503.

Bates D, Mächler M, Bolker B, Walker S. 2015. Fitting Linear Mixed-Effects Models Using lme4. Journal of Statistical Software; Vol 1, Issue 1 (2015).

Brandvain Y, Kenney AM, Flagel L, Coop G, Sweigart AL. 2014. Speciation and Introgression between Mimulus nasutus and Mimulus guttatus. Plos Genetics 10:e1004410.

Camacho C, Coulouris G, Avagyan V, Ma N, Papadopoulos J, Bealer K, Madden TL. 2009. BLAST+: architecture and applications. BMC Bioinformatics 10:421.

Cariou M, Duret L, Charlat S. 2016. How and how much does RAD-seq bias genetic diversity estimates? BMC Evolutionary Biology 16:1–8.

Chen J, Luo M, Li S, Tao M, Ye X, Duan W, Zhang C, Qin Q, Xiao J, Liu S. 2018. A comparative study of distant hybridization in plants and animals. Science China Life Sciences 61:285–309.

Coughlan JM, Matute DR. 2020. The importance of intrinsic postzygotic barriers throughout the speciation process. Philosophical Transactions of the Royal Society of London. Series B: Biological Sciences 375:20190533.

Coyne JA, Orr HA. 1989. Patterns of Speciation in Drosophila. Evolution 43:362–381.

Dowling TE, Secor CL. 1997. The Role of Hybridization and Introgression in the Diversification of Animals. Annual Review of Ecology and Systematics 28:593–619.

Durand EY, Patterson N, Reich D, Slatkin M. 2011. Testing for ancient admixture between closely related populations. Molecular Biology and Evolution.

Edelman NB, Frandsen PB, Miyagi M, Clavijo B, Davey J, Dikow RB, Garcia-Accinelli G, Van Belleghem SM, Patterson N, Neafsey DE, et al. 2019. Genomic architecture and introgression shape a butterfly radiation. Science 366:594–599.

Ellstrand NC, Prentice HC, Hancock JF. 1999. Gene Flow and Introgression from Domesticated Plants into their Wild Relatives. Annual Review of Ecology and Systematics 30:539–563.

Fitzpatrick BM. 2013. Alternative forms for genomic clines. Ecology and Evolution 3:1951–1966.

Fitzpatrick BM. 2004. Rates of evolution of hybrid inviability in birds and mammals. Evolution 58:1865–1870.

Fontaine MC, Pease JB, Steele A, Waterhouse RM, Neafsey DE, Sharakhov IV, Jiang X, Hall AB, Catteruccia F, Kakani E, et al. 2015. Mosquito genomics. Extensive introgression in a malaria vector species complex revealed by phylogenomics. Science 347:1258524.

Gautier M, Foucaud J, Gharbi K, Cézard T, Galan M, Loiseau A, Thomson M, Pudlo P, Kerdelhué C, Estoup A. 2013. Estimation of population allele frequencies from next-generation sequencing data: pool-versus individual-based genotyping. Molecular Ecology 22:3766–3779.

Gervais CE, Castric V, Ressayre A, Billiard S. 2011. Origin and Diversification Dynamics of Self-Incompatibility Haplotypes. Genetics 188:625–636.

Goldberg EE, Igić B. 2012. Tempo and mode in plant breeding system evolution. Evolution 66:3701–3709.

Grabenstein KC, Taylor SA. 2018. Breaking Barriers: Causes, Consequences, and Experimental Utility of Human-Mediated Hybridization. Trends in Ecology & Evolution 33:198–212.

Green RE, Krause J, Briggs AW, Maricic T, Stenzel U, Kircher M, Patterson N, Li H, Zhai W, Fritz MH, et al. 2010. A draft sequence of the Neandertal genome. Science 328:710–722.

Gregory TR. 2021. Animal Genome Size Database. In.

Guo Q. 2014. Plant hybridization: the role of human disturbance and biological invasion. Diversity and Distributions 20:1345–1354.

Hahn MW, Hibbins MS. 2019. A Three-Sample Test for Introgression. Molecular Biology and Evolution 36:2878–2882.

Hamlin JAP, Hibbins MS, Moyle LC. 2020. Assessing biological factors affecting postspeciation introgression. Evolution Letters 4:137–154.

Harkness A, Brandvain Y. 2020. Nonself-recognition-based self-incompatibility can alternatively promote or prevent introgression. bioRxiv:2020.2009.2029.318790.

Harris K, Nielsen R. 2016. The Genetic Cost of Neanderthal Introgression. Genetics 203:881–891.

Harrison RG, Larson EL. 2014. Hybridization, introgression, and the nature of species boundaries. Journal of Heredity 105 Suppl 1:795–809.

Heiser CB. 1973. Introgression re-examined. The Botanical Review 39:347–366.

Heiser CB. 1949. Natural hybridization with particular reference to introgression. The Botanical Review 15:645–687.

Hibbins M, Hahn M. 2021. Phylogenomic approaches to detecting and characterizing introgression. EcoEvoRxiv.

Huerta-Sánchez E, Jin X, Asan, Bianba Z, Peter BM, Vinckenbosch N, Liang Y, Yi X, He M, Somel M, et al. 2014. Altitude adaptation in Tibetans caused by introgression of Denisovan-like DNA. Nature 512:194–197.

Jagoda E, Lawson DJ, Wall JD, Lambert D, Muller C, Westaway M, Leavesley M, Capellini TD, Mirazón Lahr M, Gerbault P, et al. 2018. Disentangling Immediate Adaptive Introgression from Selection on Standing Introgressed Variation in Humans. Molecular Biology and Evolution 35:623–630.

James TY, Stajich JE, Hittinger CT, Rokas A. 2020. Toward a fully resolved fungal tree of life. Annual Review of Microbiology 74.

Jofre GI, Rosenthal GG. 2021. A narrow window for geographic cline analysis using genomic data: Effects of age, drift, and migration on error rates. Molecular Ecology Resources n/a.

Jukes TH, Cantor CR. 1969. Evolution of protein molecules. Mammalian protein metabolism 3:21–132.

Juric I, Aeschbacher S, Coop G. 2016. The Strength of Selection against Neanderthal Introgression. Plos Genetics 12:e1006340.

Kenney AM, Sweigart AL. 2016. Reproductive isolation and introgression between sympatric Mimulus species. Molecular Ecology 25:2499–2517.

Knobloch IW. 1972. Intergenic Hybridization in Flowering Plants. TAXON 21:97–103.

Korneliussen TS, Albrechtsen A, Nielsen R. 2014. ANGSD: Analysis of Next Generation Sequencing Data. BMC Bioinformatics 15:356.

Lee Y, Marsden CD, Norris LC, Collier TC, Main BJ, Fofana A, Cornel AJ, Lanzaro GC. 2013. Spatiotemporal dynamics of gene flow and hybrid fitness between the M and S forms of the malaria mosquito, Anopheles gambiae. Proceedings of the National Academy of Sciences of the United States of America 110:19854–19859.

Lenth RV. 2020. emmeans: Estimated Marginal Means, aka Least-Squares Means. Version 1.5.3.

Lewontin R, Birch L. 1966. Hybridization as a source of variation for adaptation to new environments. Evolution:315–336.

Li Y, Steenwyk JL, Chang Y, Wang Y, James TY, Stajich JE, Spatafora JW, Groenewald M, Dunn CW, Hittinger CT, et al. 2021. A genome-scale phylogeny of the kingdom Fungi. Current Biology 31:1653-1665.e1655.

Liang M, Nielsen R. 2014. The Lengths of Admixture Tracts. Genetics 197:953–967.

Lowry DB, Hoban S, Kelley JL, Lotterhos KE, Reed LK, Antolin MF, Storfer A. 2017. Breaking RAD: an evaluation of the utility of restriction site-associated DNA sequencing for genome scans of adaptation. Molecular Ecology Resources 17:142–152.

Malinsky M, Challis RJ, Tyers AM, Schiffels S, Terai Y, Ngatunga BP, Miska EA, Durbin R, Genner MJ, Turner GF. 2015. Genomic islands of speciation separate cichlid ecomorphs in an East African crater lake. Science 350:1493–1498.

Malinsky M, Matschiner M, Svardal H. 2021. Dsuite - Fast D-statistics and related admixture evidence from VCF files. Molecular Ecology Resources 21:584–595.

Malinsky M, Svardal H, Tyers AM, Miska EA, Genner MJ, Turner GF, Durbin R. 2018. Whole-genome sequences of Malawi cichlids reveal multiple radiations interconnected by gene flow. Nature Ecology & Evolution 2:1940–1955.

Mallet J. 2005. Hybridization as an invasion of the genome. Trends in Ecology \& Evolution 20:229--237.

Mallet J, Besansky N, Hahn MW. 2016. How reticulated are species? Bioessays 38:140–149.

Martin SH, Amos W. 2021. Signatures of introgression across the allele frequency spectrum. Molecular Biology and Evolution 38:716–726.

Martin SH, Davey JW, Jiggins CD. 2015. Evaluating the Use of ABBA–BABA Statistics to Locate Introgressed Loci. Molecular Biology and Evolution 32:244–257.

Martin SH, Jiggins CD. 2017. Interpreting the genomic landscape of introgression. Current Opinion in Genetics and Development 47:69–74.

Martin SH, Van Belleghem SM. 2017. Exploring Evolutionary Relationships Across the Genome Using Topology Weighting. Genetics 206:429–438.

Matute DR, Cooper BS. 2021. Comparative studies on speciation: 30 years since Coyne and Orr. Evolution 75:764–778.

Matute DR, Sepúlveda VE. 2019. Fungal species boundaries in the genomics era. Fungal Genetics and Biology 131:103249.

Miles A. 2020. cggh/scikit-allel: v1. 3.2. In: Zenodo.

Mitchell N, Campbell LG, Ahern JR, Paine KC, Giroldo AB, Whitney KD. 2019. Correlates of hybridization in plants. Evolution Letters 3:570–585.

Muirhead CA, Presgraves DC. 2016. Hybrid Incompatibilities, Local Adaptation, and the Genomic Distribution of Natural Introgression between Species. The American Naturalist 187:249–261.

Nelson RR. 1963. Interspecific Hybridization in the Fungi. Mycologia 55:104–123.

Norris LC, Main BJ, Lee Y, Collier TC, Fofana A, Cornel AJ, Lanzaro GC. 2015. Adaptive introgression in an African malaria mosquito coincident with the increased usage of insecticide-treated bed nets. Proceedings of the National Academy of Sciences of the United States of America 112:815–820.

Nosil P. 2013. Degree of symaptry affects reinforcement in Drosophila. Evolution 67:868–872.

Ortego J, Gugger PF, Sork VL. 2017. Impacts of human-induced environmental disturbances on hybridization between two ecologically differentiated Californian oak species. New Phytologist 213:942–955.

Ottenburghs J. 2021. The genic view of hybridization in the Anthropocene. Evolutionary Applications n/a.

Paradis E, Schliep K. 2019. ape 5. 0: an environment for modern phylogenetics and evolutionary analyses in R. Bioinformatics 35:526–528.

Payseur BA, Rieseberg LH. 2016. A genomic perspective on hybridization and speciation. Molecular Ecology 25:2337–2360.

Pease JB, Hahn MW. 2015. Detection and polarization of introgression in a five-taxon phylogeny. Systematic Biology 64:651–662.

Petr M, Pääbo S, Kelso J, Vernot B. 2019. Limits of long-term selection against Neandertal introgression. Proceedings of the National Academy of Sciences 116:1639–1644.

Pfeifer B, Alachiotis N, Pavlidis P, Schimek MG. 2020. Genome scans for selection and introgression based on k-nearest neighbour techniques. Molecular Ecology Resources 20:1597–1609.

Pfeifer B, Kapan DD. 2019. Estimates of introgression as a function of pairwise distances. BMC Bioinformatics 20:207.

Pickrell JK, Pritchard JK. 2012. Inference of population splits and mixtures from genome-wide allele frequency data. Plos Genetics 8:e1002967.

Prager EM, Wilson AC. 1975. Slow evolutionary loss of the potential for interspecific hybridization in birds: a manifestation of slow regulatory evolution. Proceedings of the National Academy of Sciences of the United States of America 72:200–204.

Pritchard JK, Stephens M, Donnelly P. 2000. Inference of population structure using multilocus genotype data. Genetics 155:945–959.

Prum RO, Berv JS, Dornburg A, Field DJ, Townsend JP, Lemmon EM, Lemmon AR. 2015. A comprehensive phylogeny of birds (Aves) using targeted next-generation DNA sequencing. Nature 526:569–573.

Puritz JB, Matz MV, Toonen RJ, Weber JN, Bolnick DI, Bird CE. 2014. Demystifying the RAD fad. Molecular Ecology 23:5937–5942.

Putman AI, Carbone I. 2014. Challenges in analysis and interpretation of microsatellite data for population genetic studies. Ecology and Evolution 4:4399–4428.

Rabosky DL. 2009. Ecological limits and diversification rate: alternative paradigms to explain the variation in species richness among clades and regions. Ecology Letters 12:735–743.

Rabosky DL, Santini F, Eastman J, Smith SA, Sidlauskas B, Chang J, Alfaro ME. 2013. Rates of speciation and morphological evolution are correlated across the largest vertebrate radiation. Nature Communications 4:1–8.

Racimo F, Sankararaman S, Nielsen R, Huerta-Sanchez E. 2015. Evidence for archaic adaptive introgression in humans. Nat Rev Genet 16:359–371.

Raj A, Stephens M, Pritchard JK. 2014. fastSTRUCTURE: variational inference of population structure in large SNP data sets. Genetics 197:573–589.

Reich D, Thangaraj K, Patterson N, Price AL, Singh L. 2009. Reconstructing Indian population history. Nature 461:489–494.

Rieseberg LH. 1997. Hybrid Origins of Plant Species. Annual Review of Ecology and Systematics 28:359–389.

Rieseberg LH, Wendel JF. 1993. Introgression and its consequences in plants. In. Hybrid zones and the evolutionary process. p. 109.

Roda F, Hopkins R. 2019. Correlated evolution of self and interspecific incompatibility across the range of a Texas wildflower. New Phytologist 221:553–564.

Rosenzweig BK, Pease JB, Besansky NJ, Hahn MW. 2016. Powerful methods for detecting introgressed regions from population genomic data. Molecular Ecology 25:2387–2397.

Roux C, Fraisse C, Romiguier J, Anciaux Y, Galtier N, Bierne N. 2016. Shedding Light on the Grey Zone of Speciation along a Continuum of Genomic Divergence. PLoS Biology 14:e2000234.

Sankararaman S, Mallick S, Dannemann M, Prufer K, Kelso J, Paabo S, Patterson N, Reich D. 2014. The genomic landscape of Neanderthal ancestry in present-day humans. Nature 507:354–357.

Satokangas I, Martin SH, Helanterä H, Saramäki J, Kulmuni J. 2020. Multi-locus interactions and the build-up of reproductive isolation. Philosophical Transactions of the Royal Society B: Biological Sciences 375:20190543.

Scargle JD. 1999. Publication bias (the” file-drawer problem”) in scientific inference. arXiv preprint physics/9909033.

Schluter D, Pennell MW. 2017. Speciation gradients and the distribution of biodiversity. Nature 546:48–55.

Schwenk K, Brede N, Streit B. 2008. Introduction. Extent, processes and evolutionary impact of interspecific hybridization in animals. Philosophical Transactions of the Royal Society of London. Series B: Biological Sciences 363:2805–2811.

Seehausen O. 2004. Hybridization and adaptive radiation. Trends in Ecology & Evolution 19:198–207.

Shchur V, Svedberg J, Medina P, Corbett-Detig R, Nielsen R. 2020. On the Distribution of Tract Lengths During Adaptive Introgression. G3 Genes|Genomes|Genetics 10:3663–3673.

Sievers F, Higgins DG. 2018. Clustal Omega for making accurate alignments of many protein sequences. Protein Science 27:135–145.

Small ST, Labbé F, Lobo NF, Koekemoer LL, Sikaala CH, Neafsey DE, Hahn MW, Fontaine MC, Besansky NJ. 2020. Radiation with reticulation marks the origin of a major malaria vector. Proceedings of the National Academy of Sciences 117:31583–31590.

Smith J, Kronforst MR. 2013. Do Heliconius butterfly species exchange mimicry alleles? Biology Letters 9:20130503.

Stajich JE, Harris T, Brunk BP, Brestelli J, Fischer S, Harb OS, Kissinger JC, Li W, Nayak V, Pinney DF. 2012. FungiDB: an integrated functional genomics database for fungi. Nucleic Acids Research 40:D675–D681.

Staubach F, Lorenc A, Messer PW, Tang K, Petrov DA, Tautz D. 2012. Genome Patterns of Selection and Introgression of Haplotypes in Natural Populations of the House Mouse (Mus musculus). Plos Genetics 8:e1002891.

Steensels J, Gallone B, Verstrepen KJ. 2021. Interspecific hybridization as a driver of fungal evolution and adaptation. Nature Reviews Microbiology.

Suarez-Gonzalez A, Lexer C, Cronk QCB. 2018. Adaptive introgression: a plant perspective. Biology Letters 14.

Suvorov A, Kim BY, Wang J, Armstrong EE, Peede D, D’Agostino ER, Price DK, Wadell P, Lang M, Courtier-Orgogozo V. 2021. Widespread introgression across a phylogeny of 155 Drosophila genomes. BioRxiv:2020.2012. 2014.422758.

Svedberg J, Shchur V, Reinman S, Nielsen R, Corbett-Detig R. 2021. Inferring Adaptive Introgression Using Hidden Markov Models. Molecular Biology and Evolution 38:2152–2165.

Team RC. 2020. R: A Language and Environment for Statistical Computing. Version 4.0.1.

Todesco M, Pascual MA, Owens GL, Ostevik KL, Moyers BT, Hbner S, Heredia SM, Hahn MA, Caseys C, Bock DG, et al. 2016. Hybridization and extinction. In.

van Hengstum T, Lachmuth S, Oostermeijer JGB, den Nijs HCM, Meirmans PG, van Tienderen PH. 2012. Human-induced hybridization among congeneric endemic plants on Tenerife, Canary Islands. Plant Systematics and Evolution 298:1119–1131.

Veller C, Edelman NB, Muralidhar P, Nowak MA. 2019. Recombination, variance in genetic relatedness, and selection against introgressed DNA. bioRxiv:846147.

Wangkumhang P, Hellenthal G. 2018. Statistical methods for detecting admixture. Current Opinion in Genetics and Development 53:121–127.

Whitney KD, Ahern JR, Campbell LG, Albert LP, King MS. 2010. Patterns of hybridization in plants. Perspectives in Plant Ecology, Evolution and Systematics 12:175–182.

Wiens JJ, Engstrom TN, Chippindale PT. 2006. Rapid diversification, incomplete isolation and the “speciation clock” in North American salamanders (genus Plethodon): Testing the hybrid swarm hypothesis of rapid radiation. Evolution 60:2585–2603.

Wilson AC, Maxson LR, Sarich VM. 1974. Two types of molecular evolution. Evidence from studies of interspecific hybridization. Proceedings of the National Academy of Sciences 71:2843–2847.

Zeberg H, Paabo S. 2021. A genomic region associated with protection against severe COVID-19 is inherited from Neandertals. Proceedings of the National Academy of Sciences of the United States of America 118.

Zeberg H, Paabo S. 2020. The major genetic risk factor for severe COVID-19 is inherited from Neanderthals. Nature 587:610–612.

