## Supplementary figures and images for "15 years of introgression studies: quantifying gene flow across Eukaryotes"

### Fig S2-DataMethodCorrelations

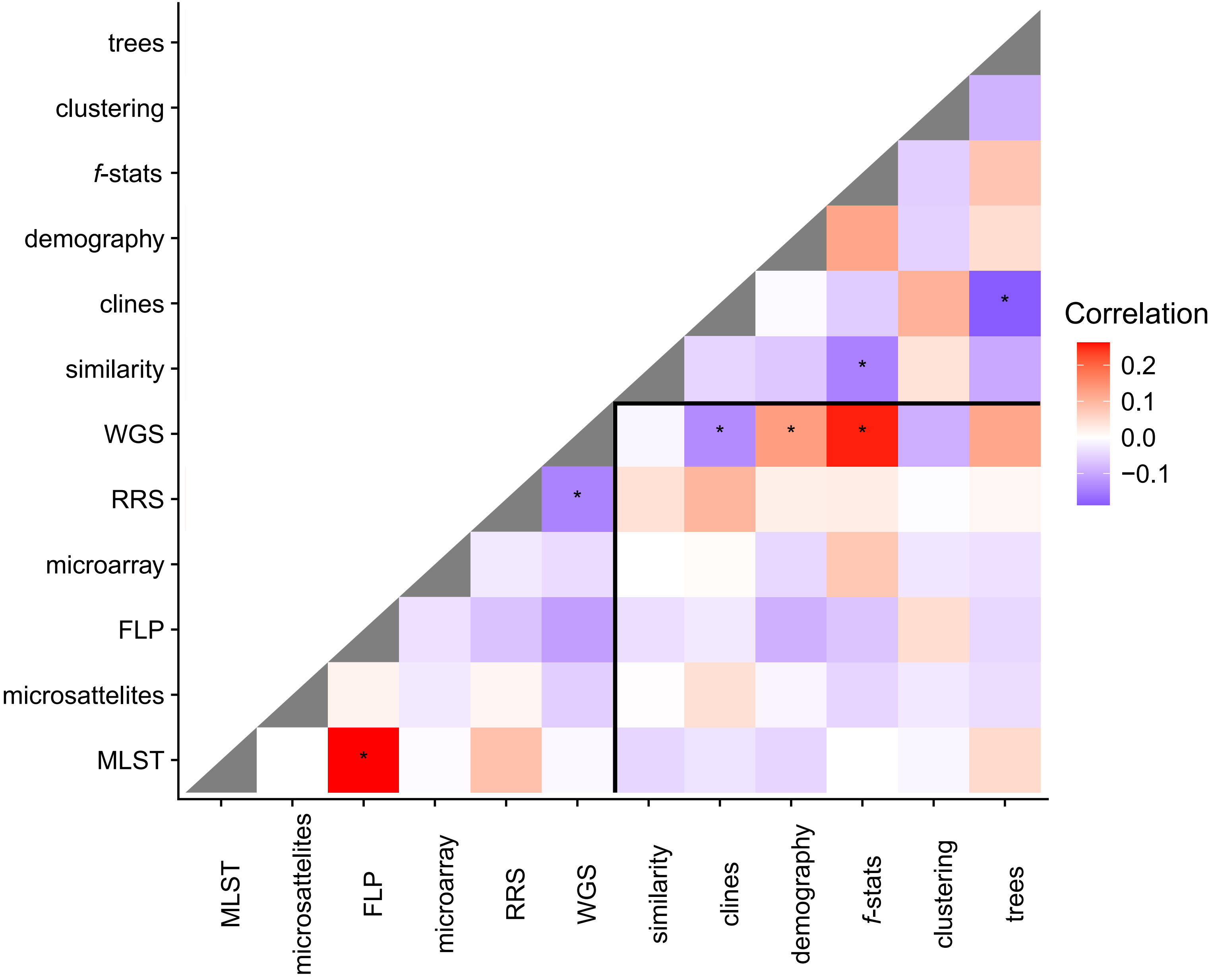

### Fig S3-TimeTree.pdf

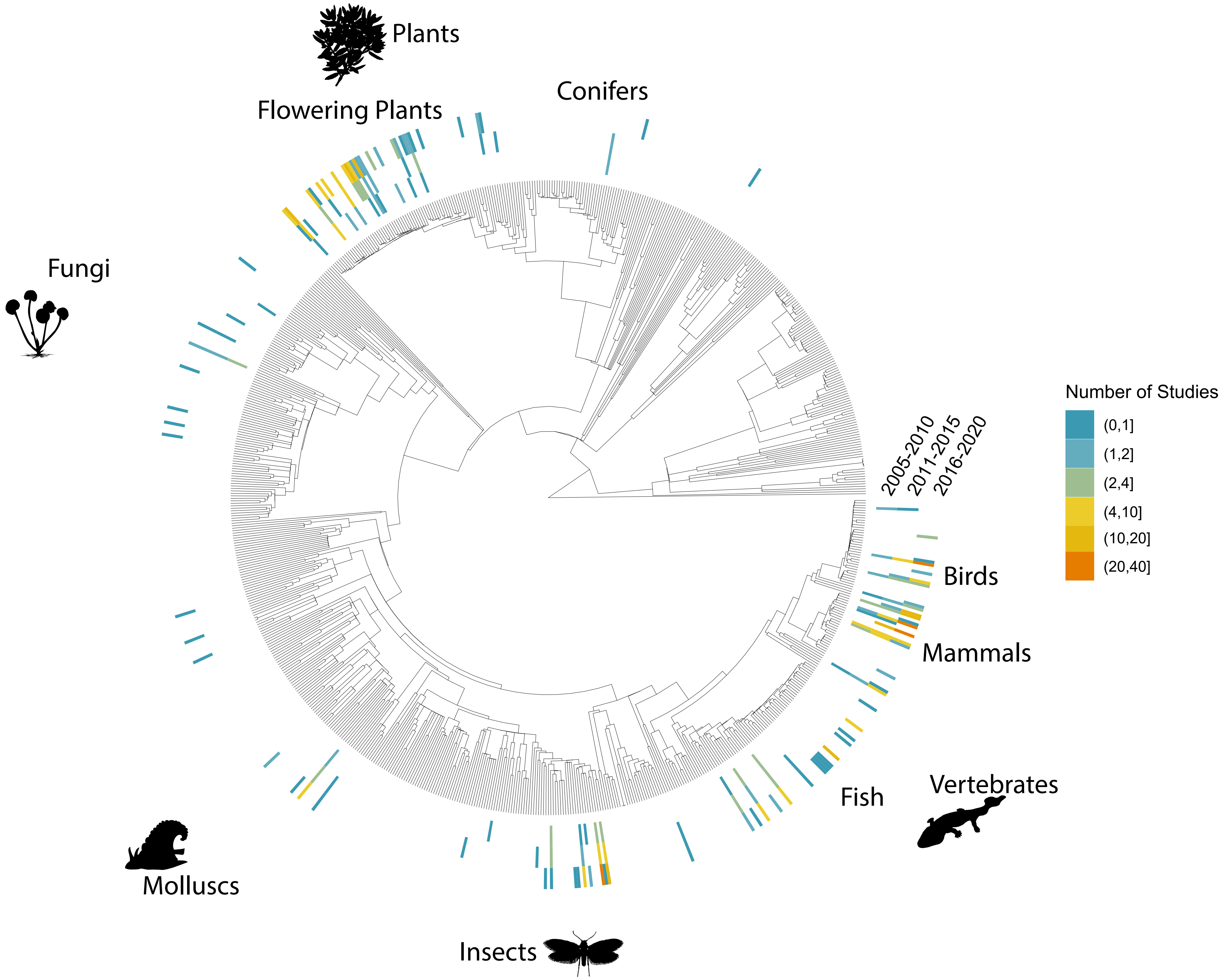

### Fig S4-DataType.pdf

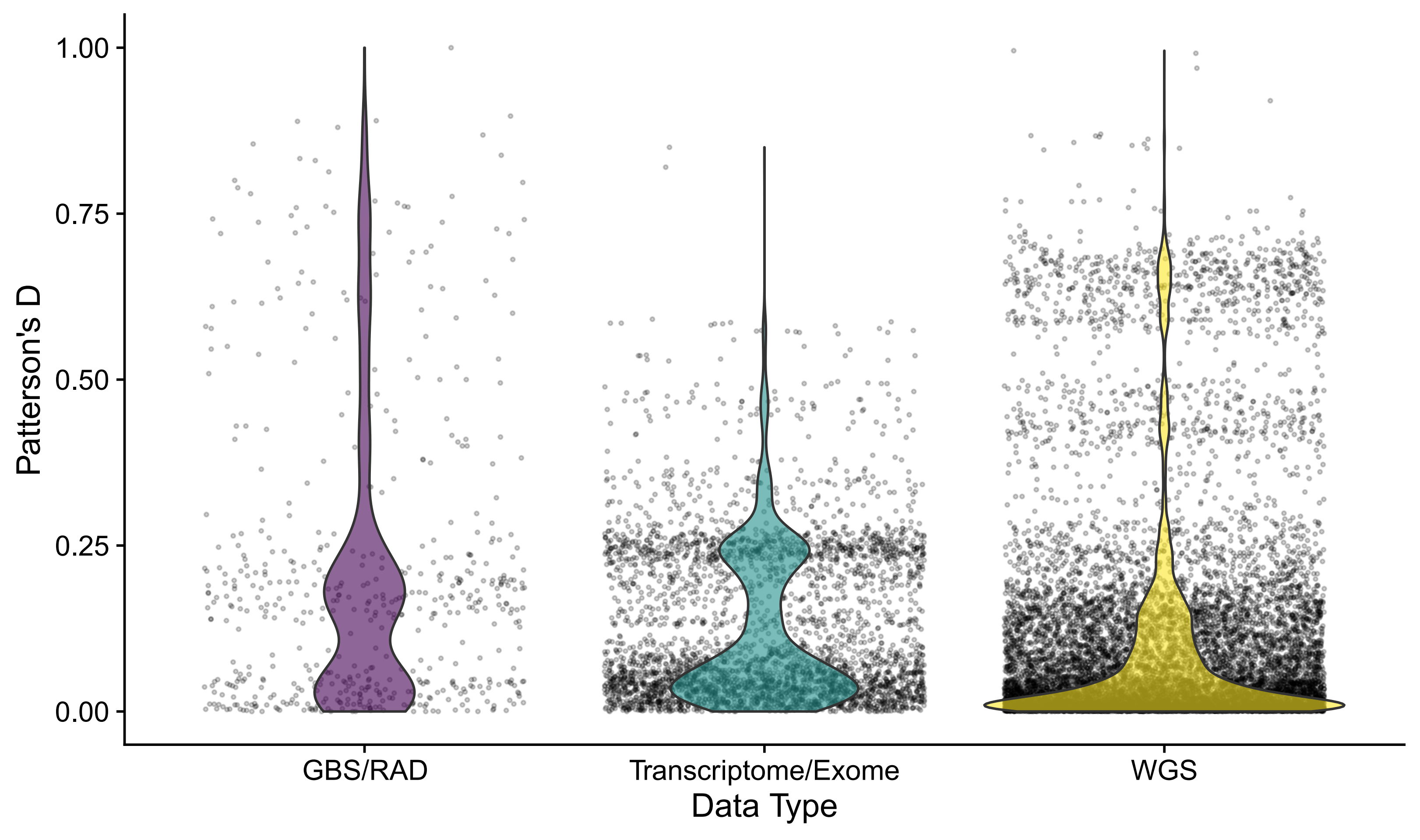

### Fig S5-ReportingType.pdf

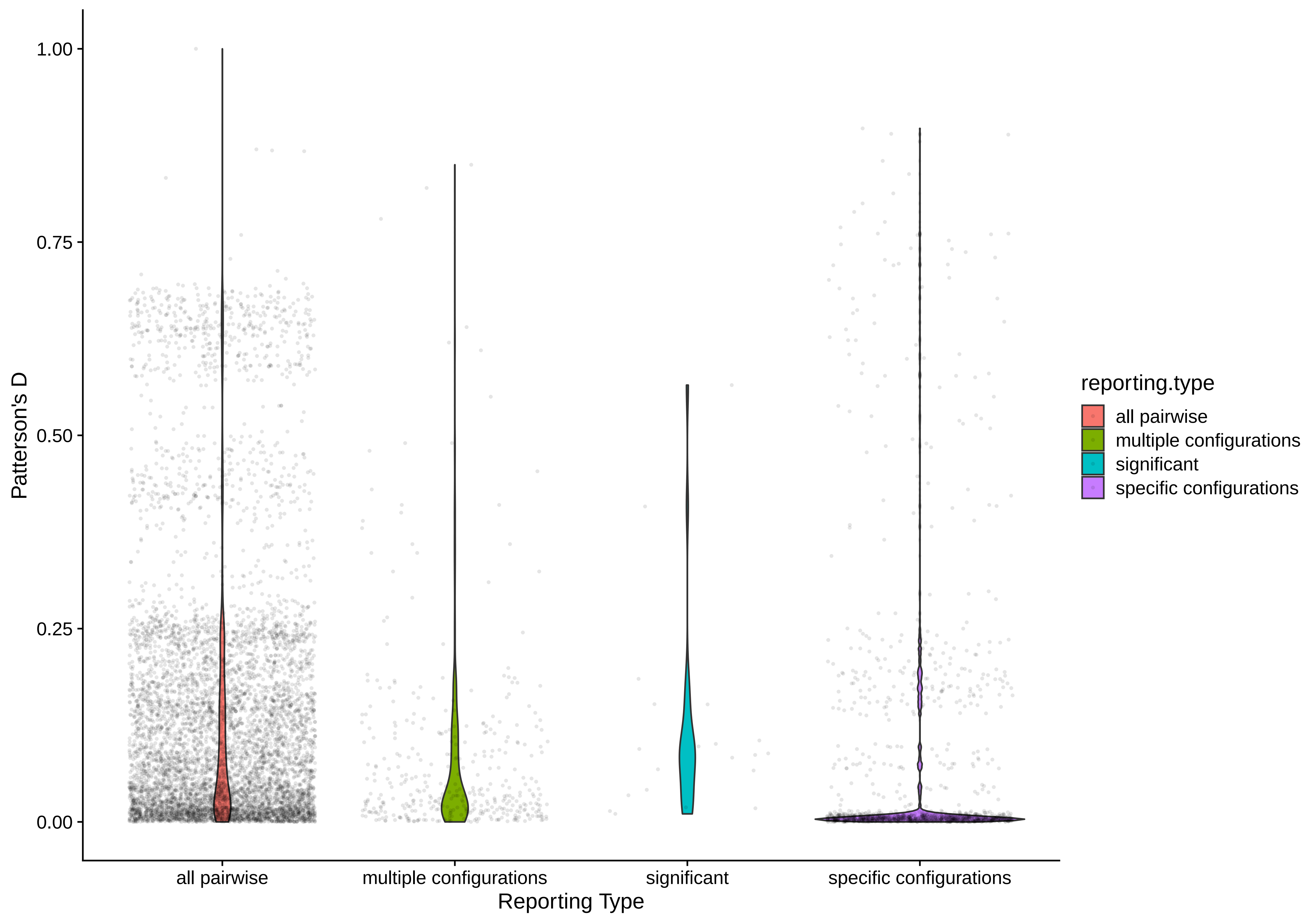

### Fig S6-HumanAssociated.pdf

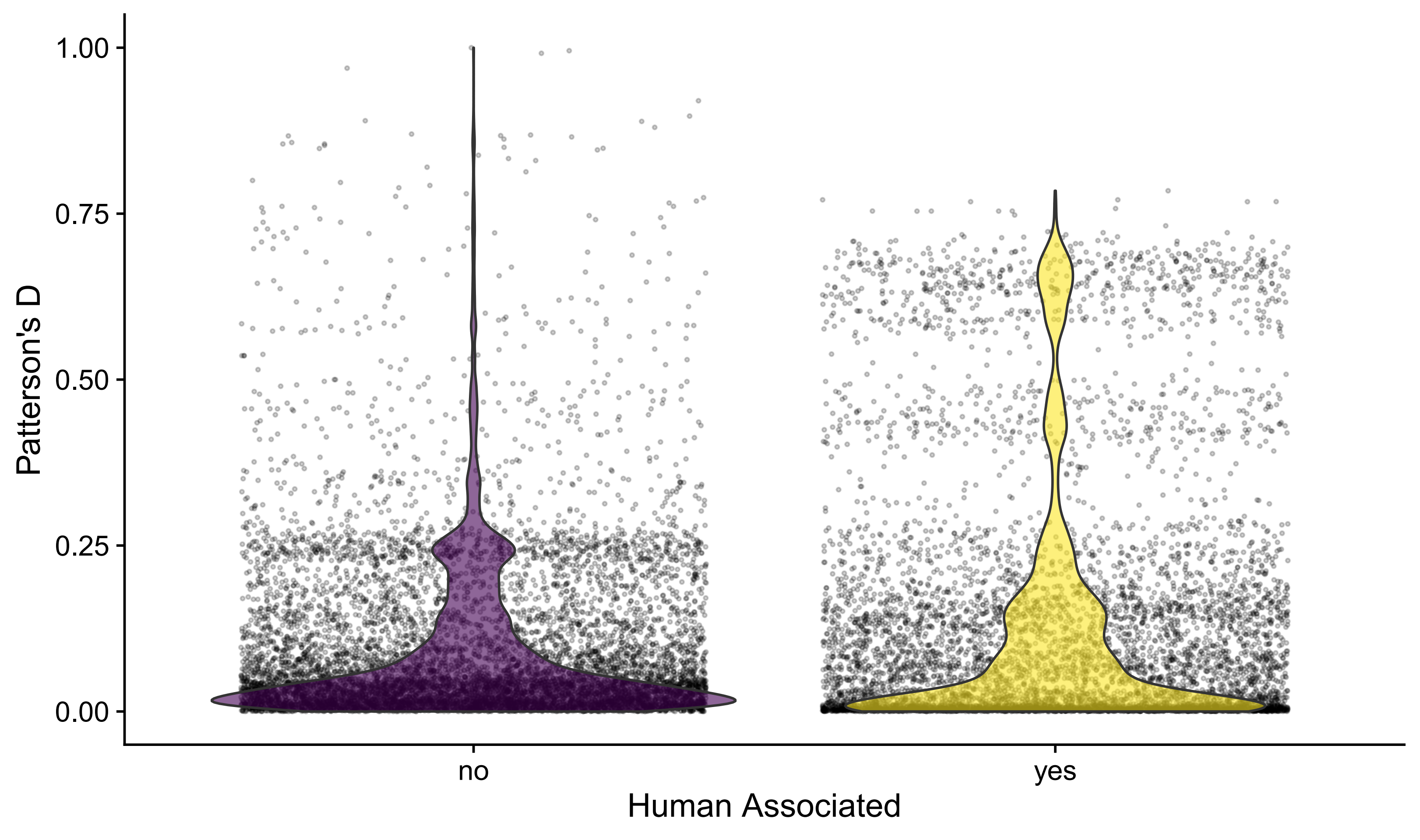

### Fig S7-SexualSystem.pdf

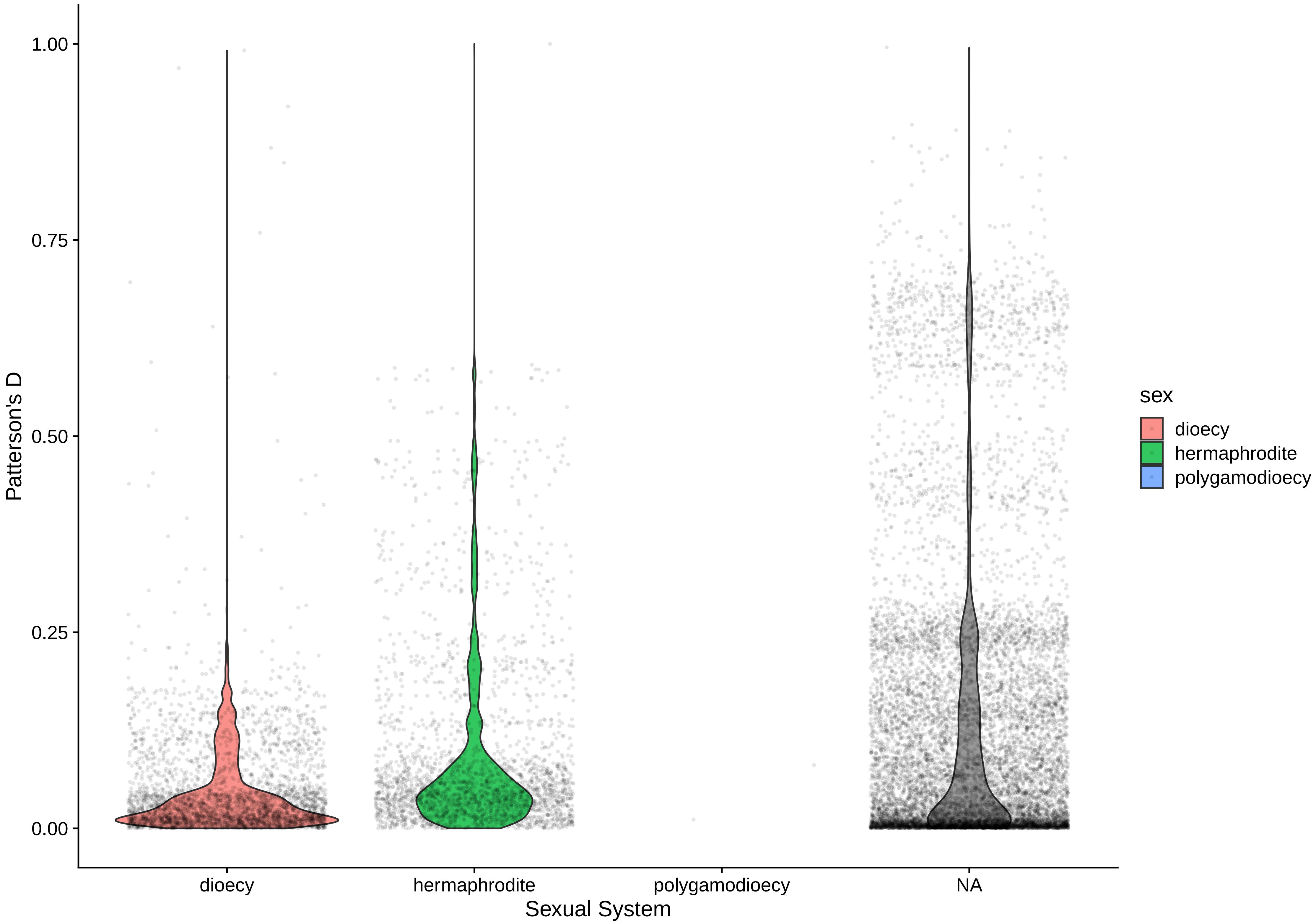

### Fig S8-Selfing_Combo.pdf

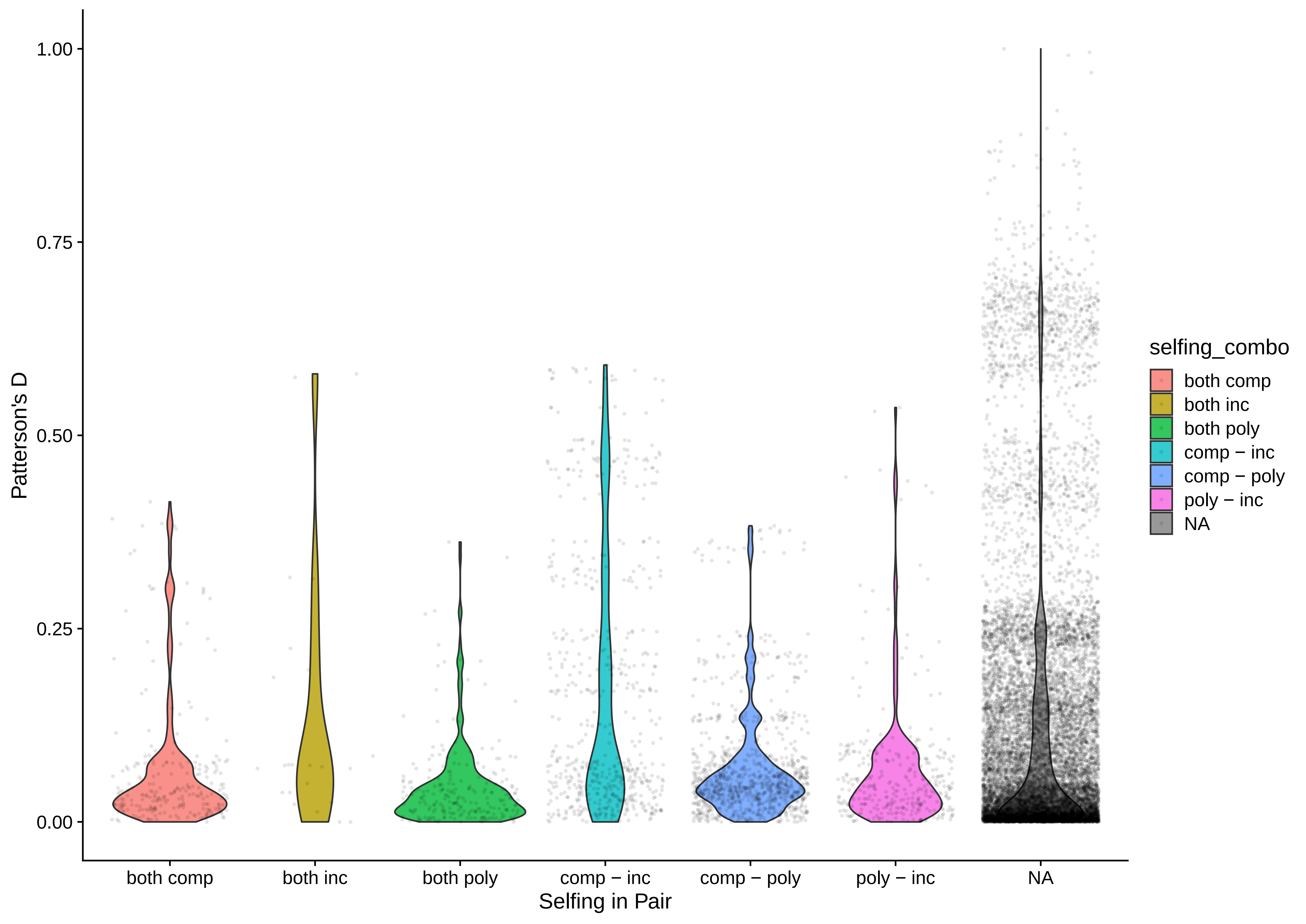
